# HSV Forms an HCMV-like Viral Assembly Center in Neuronal Cells

**DOI:** 10.1101/2020.04.22.055145

**Authors:** Shaowen White, Hiroyuki Kawano, N. Charles Harata, Richard J. Roller

**Author notes:** Corresponding Author, 3-432 BSB, 51 Newton Road, Iowa City, IA 52242.

## Abstract

Herpes simplex virus (HSV) is a neuroinvasive virus that has been used as a model organism for studying common properties of all herpesviruses. HSV induces host organelle rearrangement and forms dispersed assembly compartments in epithelial cells, which complicates the study of HSV assembly. In this study, we show that HSV forms a visually distinct unitary cytoplasmic viral assembly center (cVAC) in both cancerous and primary neuronal cells that concentrates viral structural proteins and is the site of capsid envelopment. The HSV cVAC also concentrates host membranes that are important for viral assembly, such as Golgi- and recycling endosome-derived membranes. Lastly, we show that HSV cVAC formation and/or maintenance depends on an intact microtubule network and a viral tegument protein, pUL51. Our observations suggest that the neuronal cVAC is a uniquely useful model to study common herpesvirus assembly pathways, and cell-specific pathways for membrane reorganization.

**Summary:** This study shows that HSV forms a viral assembly center in neuronal cells by reorganization of host membranes. This system is a novel and powerful tool to study herpesvirus assembly pathways and host cell membrane dynamics.

## Introduction

It is commonly observed that viruses concentrate viral and cellular components for certain processes, either final viral assembly or viral replication, at a defined subcellular compartment. In the case of retroviruses, influenza virus, rhabdoviruses, and filoviruses, the compartment of final assembly is the plasma membrane, where all virion components converge and are incorporated into the virion during viral budding (Welsch et al., 2007). In the case of reovirus, coronavirus, and hepatitis C virus (HCV), the host ER membrane is rearranged to form visually distinct viral structures that serve as either assembly factories (Tenorio et al., 2019), or replication vesicles (Knoops et al., 2008, Paul et al., 2013). It might be expected that such a concentration of virion components is especially important for the assembly of viruses that contain many diverse structural proteins and have a complex virion structure, such as herpesviruses. The virion of all herpesviruses is composed of at least three layers of structural proteins, which include a protein capsid (composed of 8 different proteins), a tegument layer (composed of as many as 23 different proteins), and an envelope containing at least 12 transmembrane proteins, which mostly are glycosylated (Loret et al., 2008). Curiously, despite the structural complexity of the mature herpesvirion, only human cytomegalovirus (HCMV), a betaherpesvirus, has been shown to form a unitary organized cVAC in infected cells (Sanchez et al., 2000).

The HCMV cVAC is a spatially organized system of membranes formed around the microtubule organization center (MTOC) (Sanchez et al., 2000), and maintenance of the cVAC is microtubule- (Indran et al., 2010) and dynein motor-dependent (Buchkovich et al., 2010). The HCMV cVAC contains enveloping capsids and concentrates various viral structural proteins and host cellular markers such as the Golgi markers TGN46, GM130, and ManII; recycling endosome marker Rab11; and early endosome marker EEA1 (Das and Pellett, 2011, Sanchez et al., 2000). The lysosome marker LAMP1 also concentrates to the HCMV cVAC but does not co-localize with viral proteins (Das and Pellett, 2011, Sanchez et al., 2000). Formation of the HCMV cVAC is dependent on the function of specific virally encoded proteins including pUL97 and pUL71. pUL97 is an early-expressing multi-functional serine/threonine kinase of which homologs exist in all known herpesviruses (Chee et al., 1989, Smith and Smith, 1989, Kato et al., 2006). pUL97, and its homolog pUL13 in HSV, are both involved in the capsid nuclear maturation and cytoplasmic assembly (Krosky et al., 2003, Azzeh et al., 2006). On the other hand, pUL71, and its homolog pUL51 in HSV, are palmitoylated membrane-associated tegument proteins that are involved in viral cytoplasmic envelopment (Womack and Shenk, 2010, Nozawa et al., 2003, Nozawa et al., 2005). Failure to express either of these proteins results in failure to form a morphologically normal cVAC, and inhibits production of infectious virus (Prichard et al., 2005, Azzeh et al., 2006, Dietz et al., 2018, Schauflinger et al., 2011).

Due to the similarity in virion structure, all herpesviruses might be expected to undergo a similar pathway for assembly, but the most extensively studied herpesvirus, HSV-1, has not been shown to form a unitary cVAC in infected cells of epithelial origin. The current understanding of HSV assembly pathway is that the viral capsid assembly and genome packaging occurs in the nucleus (Baines and Duffy, 2006), which is followed by the export of matured capsids into the cytoplasm via a well-established envelopment/de-envelopment nuclear egress process (Reynolds et al., 2002). After being released into the cytoplasm, capsids are recruited to the assembly compartment, which is dependent on tegument proteins pUL36 and pUL37 (Desai, 2000, Lee et al., 2006, Sandbaumhuter et al., 2013). Finally, the capsids bud into a vesicular compartment, resulting in double-membraned virion-containing vesicles, completing the final assembly of infectious progeny (Lv et al., 2019). It is well-known that in HSV infected cells, the microtubule network is rearranged after becoming fragmented, and the Golgi apparatus is dispersed to the periphery of the cell (Lv et al., 2019, Avitabile et al., 1995, Campadelli et al., 1993). Details of host membrane remodeling by HSV is reviewed by (Lv et al., 2019). Without a convenient centralized viral structure like the HCMV cVAC, this loss of conventional membrane structures in host cells complicates the study of HSV cytoplasmic assembly by microscopy, as mutant phenotypes tend to be ambiguous and distinguishing assembly membranes from those involved in biosynthesis and maturation of viral membrane proteins is difficult. This, in turn has made it difficult to definitively identify the viral budding site, with evidence suggesting both Golgi- and recycling endosome-derived membranes (Sugimoto et al., 2008, Hollinshead et al., 2012).

In a typical oral infection of the human host, HSV-1 initiates the infection in epithelial cells, and some of the progeny virus spreads to sensory neurons, and ascends the axons to the cell bodies in the trigeminal ganglion, where viral latency is established (Kurt-Jones et al., 2017). Replication in, and spread between neurons can lead to encephalitis in neonates and immune-compromised individuals (Whitley, 2004, Whitley, 2006), and the presence of viral components in the brain has been shown to be associated with neuron degenerative diseases, such as Alzheimer’s disease (Readhead et al., 2018). It is evident that assembly pathways of HSV in neurons and in epithelial cells are not identical, as tegument protein pUS9 is reported to be dispensable for efficient viral assembly in epithelial cells but not in neurons (DuRaine et al., 2017).

In this study, we show that HSV forms a HCMV-like unitary cVAC in mouse and human cancerous neuronal cells, as well as in primary mouse brain neurons, suggesting that the formation of a cVAC is a specific strategy for HSV assembly in neurons. Similar to the HCMV cVAC, the HSV neuronal cVAC is formed around the MTOC, remains structurally stable throughout the infection, and its maintenance depends on an intact microtubule network. The HSV cVAC also concentrates Golgi-derived membranes and recycling endosome-derived membranes, which is consistent with current HSV cytoplasmic assembly models (Lv et al., 2019). Finally, we demonstrated that HSV with a mutation in the tegument protein pUL51, a homolog of HCMV pUL71 (Roller et al., 2014), shows a disrupted cVAC phenotype correlated with a virus single-step growth defect. The functional correspondence between HCMV pUL71 and HSV-1 pUL51 suggests that the HSV-1 neuronal cVAC will prove a very valuable system for exploring the common mechanisms of herpesvirus assembly, and for exploring mechanisms for reorganization of host membrane trafficking.

## Results

### Various viral proteins concentrate at one single compartment in HSV infected neuronal cells

During studies of the function of the HSV-1 pUL51 tegument protein we observed cell-type-specific localization patterns for pUL51 and, most interestingly, organization into a unitary structure that might correspond to a cVAC in cells of the mouse neuronal CAD cell line. To determine whether this pUL51-containing structure in CAD cells might be a cVAC, we tested the localization of other viral structural proteins. CAD cells were infected with HSV-1(F) at high multiplicity of infection (m.o.i) and co-stained for various tegument and glycoproteins. As shown in Figure 1A, we observed a unitary perinuclear cluster of viral proteins in most cells (Figure 1C) in which at least six viral structural proteins including pUL11, pUL16, pUL51, gD, gE, and gL were concentrated, suggesting the viral protein cluster might be a cVAC. Note that for these viral proteins, not all of them localize to the putative cVAC at the same efficiency, for example, pUL51 almost exclusively concentrate to the cluster, while most of the gD localizes to the cytoplasmic space or cell membrane. To determine whether a similar structure forms in neuronal cells of human origin, we tested in the same way for formation of the putative cVAC in SH-SY5Y cells, a human cancerous neuronal cell line (Figure 1B). As expected, a similar putative cVAC is observed in most cells, albeit at a lower frequency (Figure 1C). In addition, unlike in CAD cells, where the putative cVAC is composed of numerous small viral protein puncta (Figure 1A), the viral protein cluster in SH-SY5Y cells typically appeared as a collection of several large granules (Figure 1B). Together, the localization pattern of viral proteins in both CAD and SH-SY5Y cells is different from it in cells of epithelial origin, such as Vero cells (Figure 1D), suggesting that the mode of viral component trafficking is cell context-dependent.

**Figure 1.**
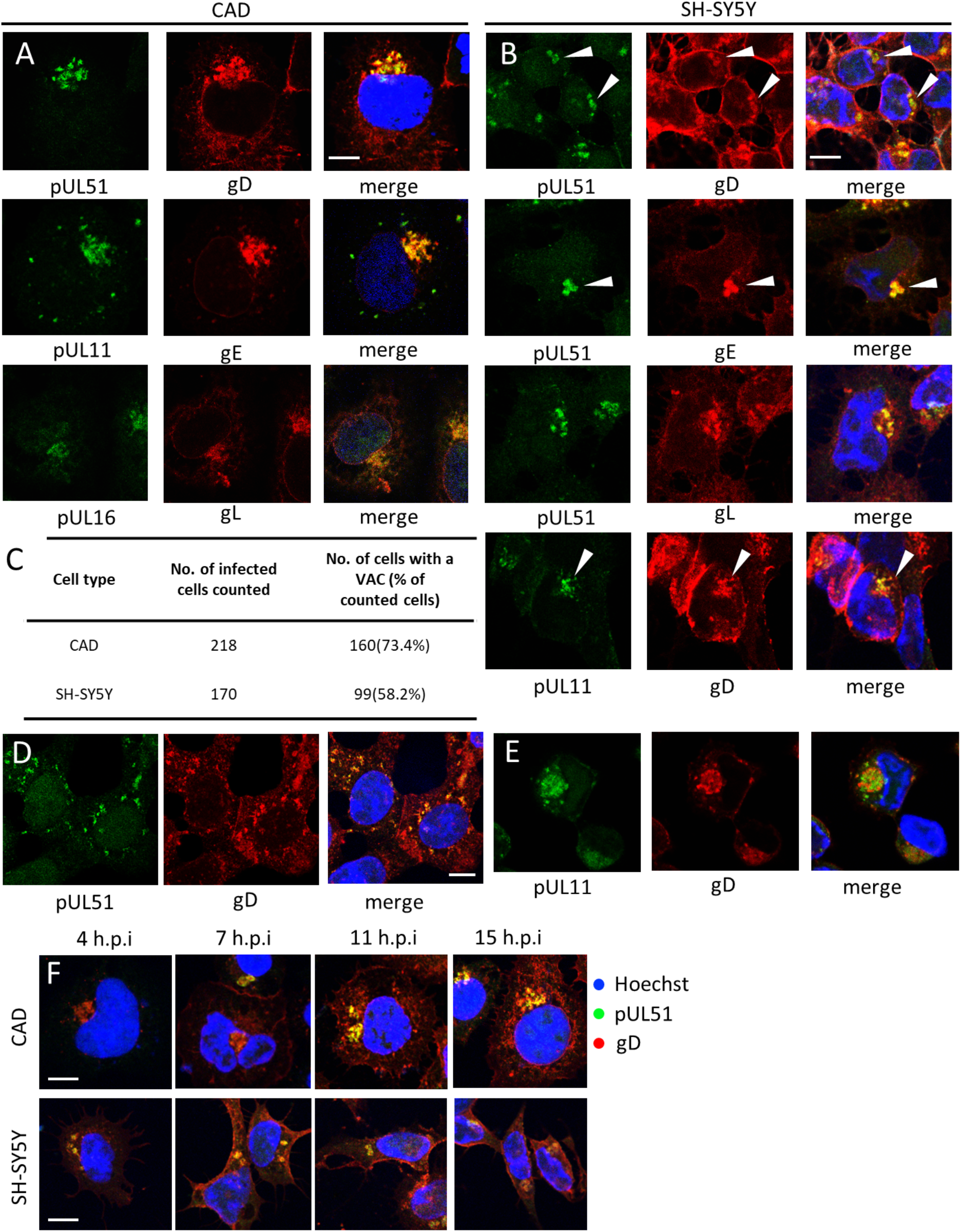
HSV forms a unitary viral protein concentration in infected cancerous neuronal cells. CAD (A) and SH-SY5Y (B) cells were infected with WT HSV-1(F) virus at m.o.i=5 and fixed at 12 h.p.i before staining for indicated viral proteins with antibodies. White arrow heads indicate representative concentrations of viral proteins. (C) Quantification of the percentage of cells that form a viral protein concentration, which was defined by a unitary concentration of co-localizing gD and pUL51 puncta on one side of the nucleus. (D) Viral protein puncta were dispersed in infected Vero cells. Vero cells were infected with WT HSV-1(F) virus at m.o.i=5 and fixed at 12 h.p.i. (E) CAD cells were infected with HSV-2 strain R519 BAC virus at m.o.i=5 and fixed at 12 h.p.i. (F) Time course of the HSV unitary viral protein concentration. CAD and SH-SY5Y cells were infected as in (A) and fixed at indicated time points before stained for pUL51 (green signal) and gD (red signal). Overlay images were chosen to represent the population of infected cells at each time point. Scale bars in all panels represent 10 μm.

To examine whether the formation of the putative cVAC is a specific property of HSV-1, we infected CAD cells with an HSV-2 bacterial artificial chromosome (BAC)-derived virus (Figure 1E). As expected, a cluster of viral proteins was observed in infected cells, although pUL11 and gD did not completely colocalize with each other within the cluster (Figure 1E).

To characterize the temporal dynamics of the HSV-1 putative cVAC, CAD and SH-SY5Y cell were subjected to synchronized infection and fixed at various time points after infection. As shown in Figure 1F, the putative cVAC was formed as early as 4 hour-post-infection (h.p.i) and was maintained as late as 15 h.p.i, and its gross morphology showed no obvious change despite the progression of the cytopathic effect (CPE) caused by the infection. Further time points are not shown due to severe cytoplasmic space shrinkage associated with cell rounding. The stability of the putative cVAC suggests that it is not an intermediate stage of CPE progression.

Immortalized cell lines often differ from their primary counterparts in many ways. To determine whether formation of a putative cVAC is a peculiarity of immortalized neuronal cells, primary mouse cortical neurons were tested. Cortical neurons from newborn mouse pups were harvested as described in Material and Methods and allowed to differentiate *in vitro* for 3 days on a layer of primary mouse astrocyte feeder cells. The mixed culture was then infected with HSV-1(F) and fixed 12 h.p.i before being subjected to immunofluorescent microscopy. Interestingly, a putative cVAC was formed in cells that were stained positive for neuron marker MAP2 (Figure 2, B and C, white arrows), while in astrocytes, viral proteins form dispersed puncta that resemble the viral protein distribution in epithelial cells (Figure 2C, cyan arrow). These results suggest that the formation of a putative cVAC is a property of both primary and immortalized neuronal cells, and may be specific for cells of the neuronal lineage.

**Figure 2.**
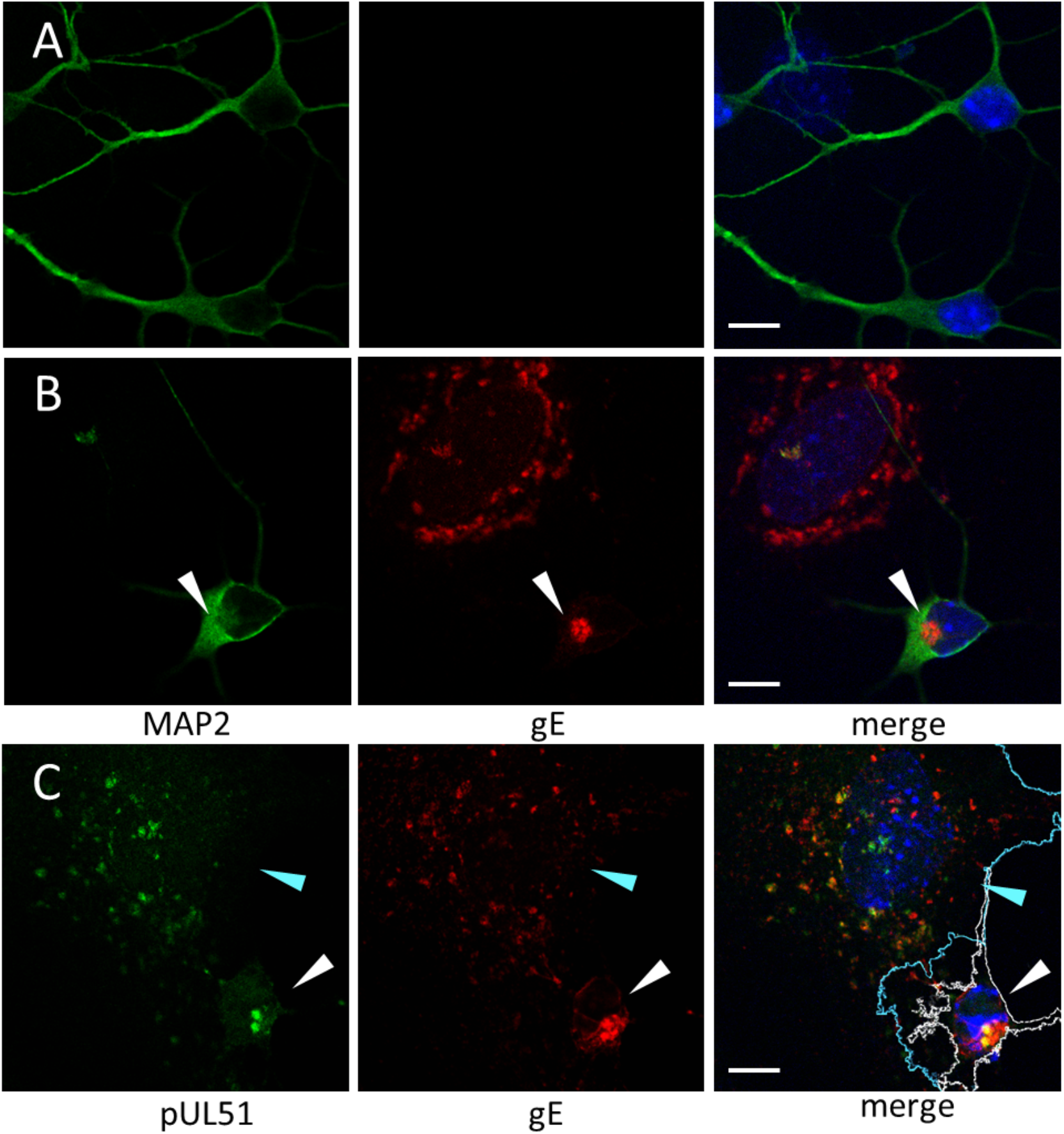
Formation of viral protein concentration in infected primary mouse neurons. Mouse neurons and mouse astrocyte feeder cells mixed cultures were mock-infected (A) or infected with WT HSV-1(F) virus (B-C) for 11 hs before fixation and staining. (A-B) Cells were stained for gE (red) and neuron marker MAP2 (green). Small nuclei with intense Hoechst stain (blue) represent mouse neurons. Large nuclei with faint Hoechst stain represent mouse astrocytes. The viral protein concentration at the cell body of the infected neuron is indicated with a white arrowhead. (C) Infected cells were stained for gE (red) and pUL51 (green). In merged image, the infected mouse astrocyte was indicated with a cyan arrowhead and outlined with cyan lines. The infected mouse neuron was indicated with a white arrowhead and outlined with white lines. Both outlines were defined by phalloidin staining (not shown). Scale bars represent 10 μm.

### The viral protein cluster is the site of capsid envelopment

It is well-established that HCMV forms a cVAC in infected cell. The HCMV viral protein cluster is formed around the MTOC (Sanchez et al., 2000) and is the site of capsid envelopment (Gilloteaux and Nassiri, 2000). For most cell types, the MTOC is formed around centrosome, which can be defined by staining for a centrosome-specific microtubule protein γ-tubulin (Sanchez and Feldman, 2017). To test the hypothesis that the cluster of viral proteins that were observed in HSV-infected neuronal cells is an HCMV cVAC-like viral assembly compartment, the distribution of viral capsids in relationship to the viral protein cluster and centrosome was investigated. First, CAD cells were infected with a recombinant BAC-derived HSV-1 in which the minor capsid protein pUL35 (VP26) was fused with a red fluorescent protein (RFP), and then the localizations of cytoplasmic capsids, viral tegument protein pUL11, and γ-tubulin were determined. As shown in Figure 3A, most cytoplasmic capsids were concentrated within the “clouds” of pUL11 staining, which surrounded the centrosome, suggesting that the viral protein cluster (putative cVAC) is a viral assembly compartment.

**Figure 3.**
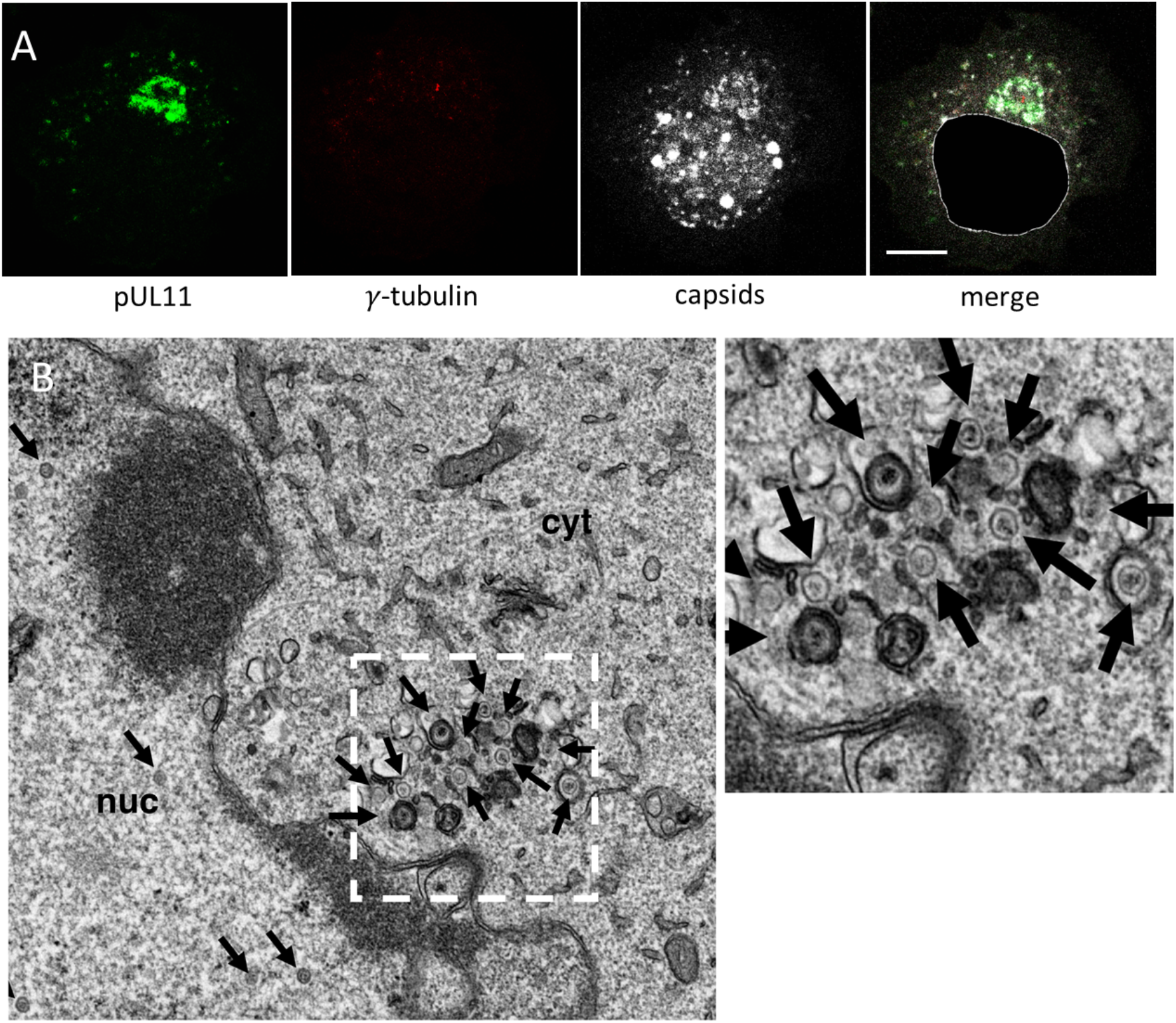
Concentration of capsids and viral proteins around the centrosome in infected CAD cells. (A) CAD cells were infected with HSV-1(F) UL35-RFP BAC virus at m.o.i=5 before fixation at 12 h.p.i. Infected cells were stained for pUL11 (green) and γ-tubulin (infrared fluorescent signal, shown in red). RFP-capsid is shown in white. In the merged image, the nucleus of the cell was masked based on Hoechst stain. The scale bar represents 10 μm. (B) CAD cells were infected with WT HSV-1(F) BAC virus at m.o.i=5 for 16 hs before fixation and TEM analysis. Black arrows indicate various capsids in the cell. The inset on the right shows zoomed region outlined by white dash lines.

To confirm that capsid envelopment occurs within a unitary structure in CAD cells, transmission electron microscopy (TEM) was performed on infected CAD cells. Whenever more than ten capsids were observed in a microscopy section, most of them were concentrated at one single perinuclear compartment in the cells, in which various stages of enveloping capsids can be identified (Figure 3B). Taken together, these results strongly suggest that HSV forms a cVAC at which capsids are enveloped in infected neurons.

### HSV putative cVAC is not an autophagosome or aggresome

The double-membrane nature of some structures that were observed in TEM (Figure 3B), suggested the possibility that the putative cVAC might instead be an autophagic compartment, as autophagy is a major mechanism of anti-viral response in neuronal cells (Orvedahl et al., 2007). Autophagy leads to the formation of a double-membrane structure called autophagosome, which engulfs cellular components and intracellular pathogens within itself before fusing with lysosomes for degradation (Tanida, 2011, Parzych and Klionsky, 2014). In mammalian cells, the membranes that form an autophagosome are defined by the incorporation of an autophagy marker LC3 (Parzych and Klionsky, 2014). To examine if the putative cVAC in neuronal cells contains autophagosome marker LC3, CAD and SH-SY5Y cells were transfected with plasmid expressing either a GFP-conjugated LC3 protein or GFP control and then infected with HSV-1 (Figure 4, A-D). In uninfected cells, GFP-LC3 remains mostly cytoplasmic with some localizing to membranes, forming puncta structures (Figure 4C). Upon infection, these GFP-LC3 puncta do not appear to concentrate to any subcellular compartment or co-localize with viral tegument protein pUL11 or viral glycoprotein gD, suggesting that the HSV putative cVAC is not a canonical autophagosome.

**Figure 4.**
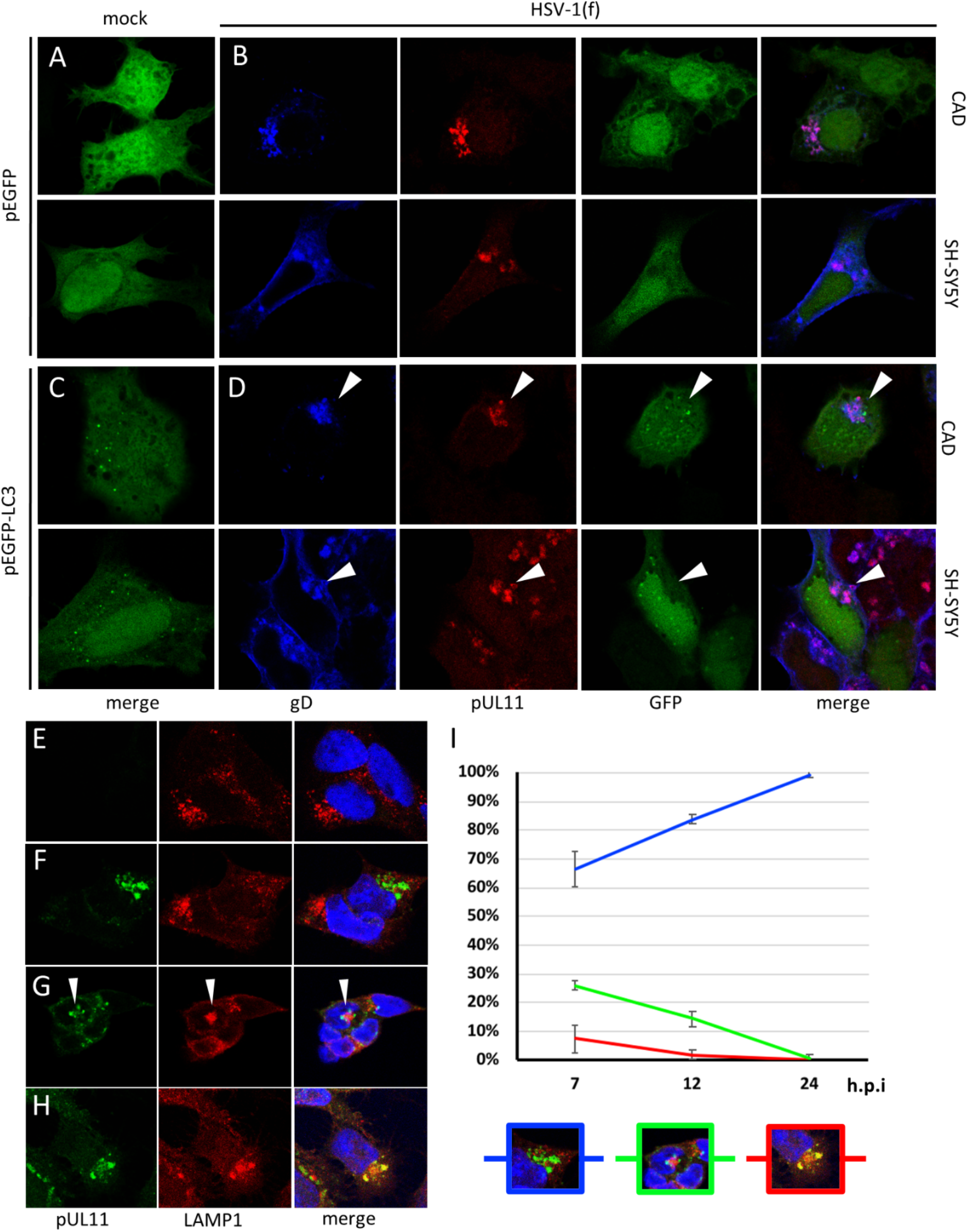
The viral protein concentration is not a canonical degradation compartment. CAD and SH-SY5Y cells were either transfected with GFP-expressing plasmid (A-B) or transfected with GFP-LC3-expressing plasmid (C-D) at least 36 hs before infections. Transfected cells were then either mock-infected (A, C) or infected with WT HSV-1(F) virus at m.o.i=5 (B, D) and fixed at 12 h.p.i. Mock-infected and infected cells were stained with anti-gD antibody (blue) and anti-UL11 antibody (red). Individual channels are only shown for infected cells. In GFP-LC3-expressing cells, the viral protein concentration is indicated with white arrowheads. (E-H) Immunofluorescent images of pUL11 (green) and LAMP1 (red) in mock-infected (E) or WT HSV-1(F)-infected (F-H) SH-SY5Y cells (m.o.i=5). The white arrowhead indicates the structure in which LAMP1 concentration is surrounded by several pUL11 puncta. (I) Time course of the percentage of each type of LAMP1 structure in infected cells (blue: F, green: G, red: H). SH-SY5Y cells were infected as in (F) but fixed at different time points. Error bars represent standard deviation of three independent experiments. For each experiment, at least 200 cells from random fields were included.

When overexpressed in cells, misfolded proteins can concentrate around the centrosome in a microtubule-dependent manner, which forms a protein aggregation termed the aggresome, with lysosomes concentrated on the outer rim (Johnston et al., 1998). To examine whether the HSV putative cVAC is instead an aggresome-like degradation compartment, LAMP1, a lysosome marker, was stained in infected SH-SY5Y cells. As shown in Figure 4F, the majority of infected cells has reduced LAMP1 positive structures at 12 h.p.i, and viral tegument protein pUL11 does not particularly co-localize or form organized spatial structures with LAMP1 puncta, suggesting that HSV putative cVAC is a *bona fide* viral assembly site. Interestingly, we also noticed that a minor population of infected cells have shown distinct LAMP1 localization patterns:(i) pUL11 puncta rimming around one large intense LAMP1 aggregate (Figure 4G), and (ii) co-localization between pUL11 and LAMP1 (Figure 4H). To examine whether these two patterns represent intermediate stages of HSV cVAC formation, infected SH-SY5Y cells were fixed at various time points, co-stained for pUL11 and LAMP1, and counted for the number of cells exhibiting each distinct pattern. The proportion of cells that showed no association between pUL11 and LAMP1 increased with time, while those that showed either of the two patterns of association decreased as infection progressed (Figure 4I), suggesting that association of the cVAC with LAMP1 represents early stages of lysosome morphology alteration by HSV infection.

### Golgi-derived membrane is concentrated at the HSV cVAC

It has been long hypothesized that Golgi-derived membrane is an important component of the HSV assembly compartment (Sugimoto et al., 2008), and Golgi markers are typically found in the vicinity of tegument protein puncta and viral capsids in HSV infected epithelial cells (Sugimoto et al., 2008, Hollinshead et al., 2012). On the other hand, in HCMV infected cells, both trans- and cis-Golgi markers are concentrated in the cVAC (Das et al., 2007). To elucidate the spatial relationship of Golgi-derived membrane and the HSV cVAC in neuronal cells, infected CAD and SH-SY5Y cells were co-stained for Golgi membrane markers GM130 (cis-Golgi) or Golgin97 (trans-Golgi) along with viral tegument protein pUL51. As shown in Figure 5, membranes positive for GM130 concentrate at the same place as pUL51 in infected cells. Interestingly, in SH-SY5Y cells, GM130 positive membranes were observed to form a tubular structure partially or completely surrounding, but not coinciding with pUL51 staining.

**Figure 5.**
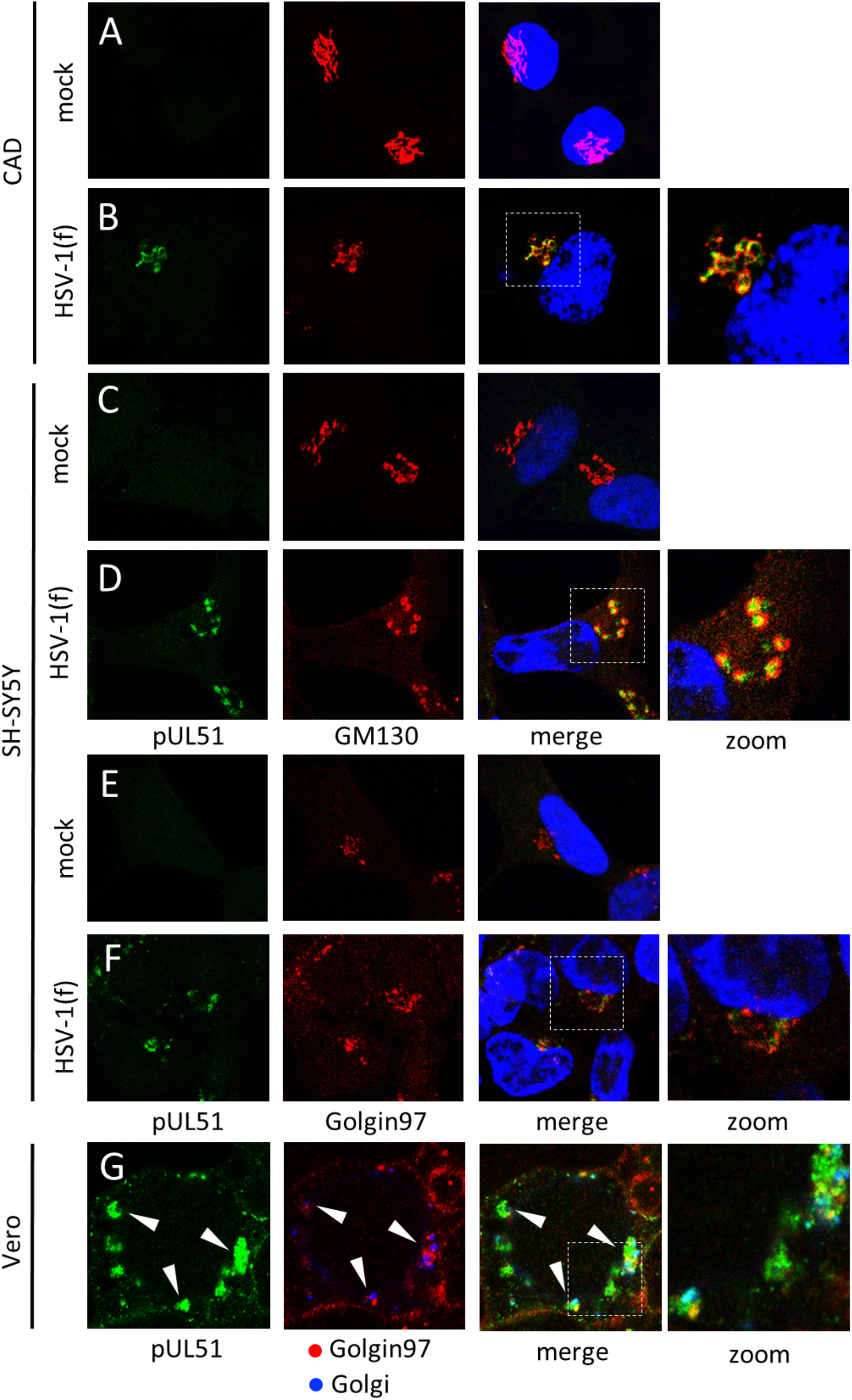
Golgi-derived membranes concentrate to the HSV cVAC. CAD and SH-SY5Y cells were mock-infected (A, C, E) or infected with WT HSV-1(F) (B, D, F) as described in Figure 1 and stained for pUL51 (green) or Golgi markers (red), including GM130, a Golgi matrix protein, and Golgin97, a trans-Golgi marker. (G) Vero cells were transduced with a baculovirus expressing a cis- and medial-Golgi marker EGFP-N-acetylgalactosaminyl transferase (rendered in blue) 24 h before HSV infection. Infected cells were then fixed 16 h.p.i. and stained for pUL51 (green) and Golgin97 (red) with respective antibodies. White arrows indicate clusters of pUL51 in adjacent to Golgi membranes. All insets to the right show the zoomed regions outlined by white dash lines in the overlay images.

In infected SH-SY5Y cell, Golgin97 positive membranes also concentrated at the same subcellular compartment as the HSV cVAC, however, individual Golgin97 puncta did not co-localize with pUL51 puncta (Figure 5F). This is consistent with our observation of infected Vero cells, in which pUL51 and Golgi membranes localized in adjacent to each other at multiple dispersed foci (Figure 5G, white arrowheads), further supporting the hypothesis that the HSV cVAC is where viral assembly occurs.

To further compare the similarity between the HSV cVAC and the HCMV cVAC, a fluorescent protein fused with an ER-localizing signal (Denecke et al., 1992) was expressed in CAD or SH-SY5Y cells to mark ER membranes before the infection. As shown in Supplemental Figure 1 (Figure S1), ER-derived membranes do not concentrate at the HSV cVAC in either infected cell line but appear to be excluded from the cVAC area.

### Rab11, but not Rab5, is concentrated at the HSV cVAC

Rab family proteins are small GTPases that guide the trafficking of various vesicles in the endosomal/secretory apparatus and contribute to the identity of different cytoplasmic membrane compartments (Stenmark, 2009). Specific Rab family proteins have been shown to play role in the assembly of alpha- and beta-herpesviruses. For example, siRNA against Rab5 or Rab11 each can result in more than 10-fold assembly defect for HSV-1 in HeLa cells (Hollinshead et al., 2012), suggesting that both are important for virus production in epithelial cells. Also, certain subsets of Rab family proteins concentrate to the cVAC in HCMV infected cells and contribute to its morphology (Das et al., 2007, Das and Pellett, 2011, Lucin et al., 2018, Indran and Britt, 2011). These observations allow us to hypothesize that the maintenance of the HSV cVAC in neuronal cells would also involve the manipulation of Rab family proteins, specifically Rab5 and Rab11. To characterize the dynamics of Rab5 and Rab11 in infected neuronal cells, uninfected or infected SH-SY5Y and CAD cells were fixed and co-stained with the antibody against gE in combination with the antibody against Rab5 or Rab11 (Figure 6). In uninfected CAD and SH-SY5Y cells, both Rab5 and Rab11 concentrate to one perinuclear region (Figure 6, A, G, D, and K), similar to the observation of others (Hollinshead et al., 2012, Salgado-Polo et al., 2020). Since it is established that Rab5 and Rab11 concentrate to the MTOC (Ullrich et al., 1996), this Rab5/11-concentration region in CAD and SH-SY5Y cells likely contains the centrosome. To show that the concentration of Rab5 and Rab11 around the centrosome is actively maintained in a microtubule-dependent manner, uninfected CAD and SH-SY5Y cells were treated with 5μg/ml of a small molecule microtubule polymerization inhibitor nocodazole. At similar concentrations, CAD and SH-SY5Y cells stayed viable under nocodazole treatment, as observed by others (Li et al., 2006, Chen et al., 2014, Ren et al., 2009) and by us (data not shown). After treatment, both Rab5 and Rab11 no longer concentrated at one single compartment (Figure 6, B, E, H, and L), suggesting Rab5 and Rab11 may be transported toward the centrosome on microtubule motors in CAD and SH-SY5Y cells.

**Figure 6.**
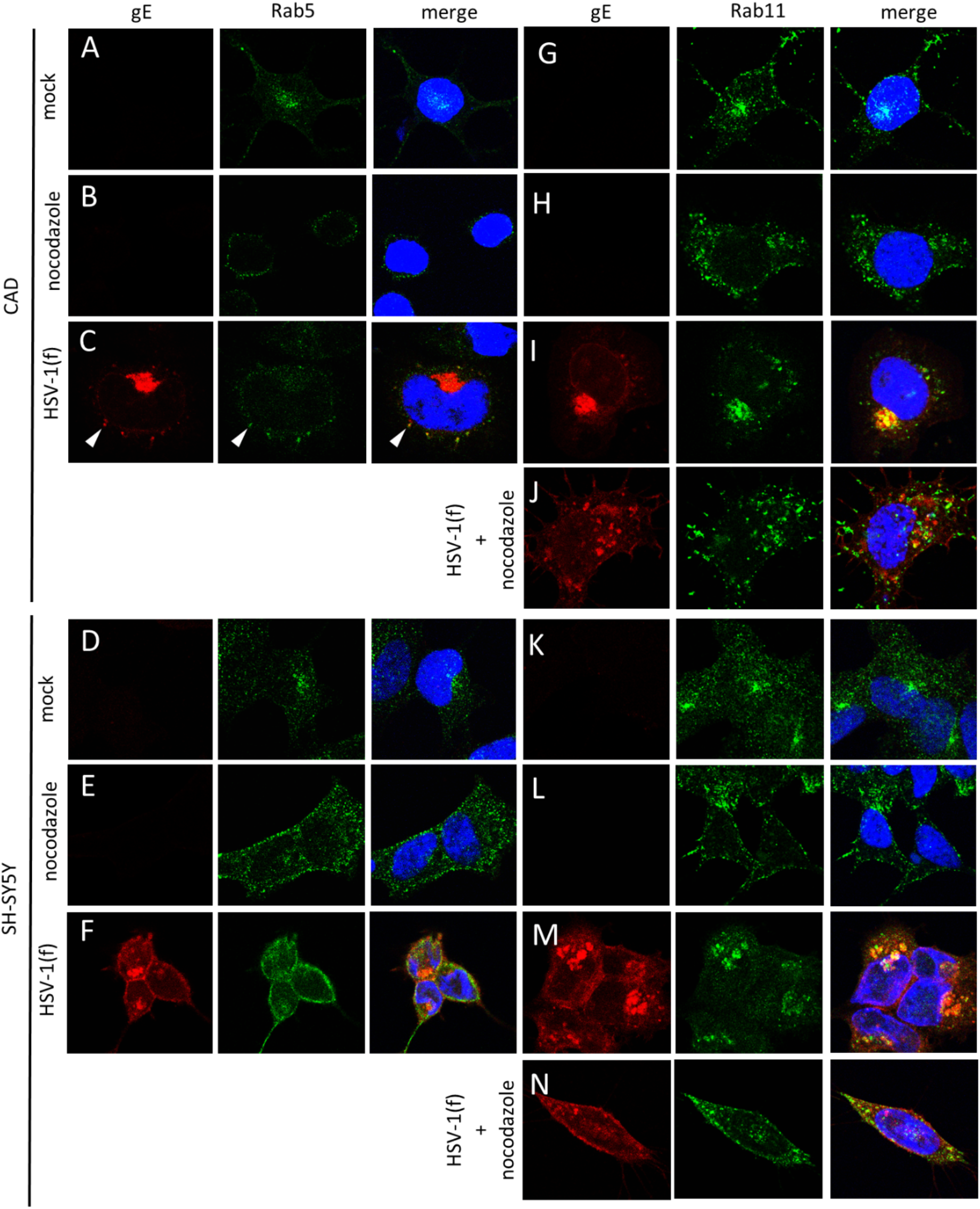
Localization of Rab5 and Rab11 in relationship to the HSV cVAC. CAD (A-J) and SH-SY5Y (D-N) cells were stained for gE (red) and a Rab family protein (Green; Rab5 or Rab11). (A, G, D, K) Mock-infected cells treated with DMSO. (B, H, E, L) Mock-infected cells treated with nocodazole for 9 hs. (C, I, F, M) WT HSV-1(F) infections (m.o.i=5) were synchronized as described in materials and methods. At 3 h.p.i, the medium was replaced with medium containing DMSO. Then, cells were fixed at 12 h.p.i. and stained. (J, N) Cells were infected as in (I, M), except at 3 h.p.i, the medium was replaced with medium containing nocodazole. White arrowheads in (C) indicate a periphery punctum that is positive for both gE and Rab5.

In both CAD and SH-SY5Y cells, infection with HSV resulted in Rab11 concentration at the cVAC defined by gE staining (Figure 6, I and M). Furthermore, in CAD cells, Rab 5 appeared to be excluded from the cVAC since no overlapping signal was observed (Figure 6C), whereas in SH-SY5Y cells, Rab5 was distributed throughout the cytoplasm, including the cVAC area (Figure 6F). Interestingly, in CAD cells, whose larger cytoplasmic space allowed clearer identification of individual puncta signals, Rab5 puncta co-localized with gE puncta that were not part of the cVAC (Figure 6C, white arrowhead).

Since Rab11 was actively transported to the centrosome in a microtubule-dependent manner in uninfected cells (Figure 6, B, E, H, and L), we hypothesize that the concentration of Rab11 to the cVAC in an HSV infection is also microtubule-dependent. To test this hypothesis, CAD and SH-SY5Y cells were treated with nocodazole three hours after synchronized infection. As shown in Figure 6, J and N, in nocodazole treated HSV infected cells, Rab11 no longer accumulates at a defined concentration; instead, Rab11 puncta disperse throughout the cell, with no obvious localization pattern, suggesting microtubule polymerization is required for Rab11 concentration to the HSV cVAC.

### Microtubule filament is essential for the maintenance of HSV cVAC

As shown in Figure 6, treatment with nocodazole not only changes the localization pattern of Rab11 puncta in HSV infected cells, but also prevents gE from concentrating to the cVAC. Since it is well-established that cVAC formation in HCMV infected cells is microtubule- and dynein motor-dependent (Indran et al., 2010, Buchkovich et al., 2010), we hypothesize that the similar subsets of cytoskeleton components are also required for HSV cVAC formation. To test whether microtubule polymerization is required for HSV cVAC formation, uninfected and synchronously infected CAD and SH-SY5Y cells were treated with 5μg/ml nocodazole. This concentration of nocodazole caused dispersion of microtubule filaments in uninfected CAD and SH-SY5Y cells (Figure 7A). In infected CAD and SH-SY5Y cells, treatment with nocodazole results in failure to form or maintain an MTOC-centered cVAC (Figure 7, C and D), and increasing concentration of nocodazole profoundly disrupted cVAC formation in a dose-dependent manner (Figure 8A), suggesting that HSV cVAC formation is microtubule-dependent.

**Figure 7.**
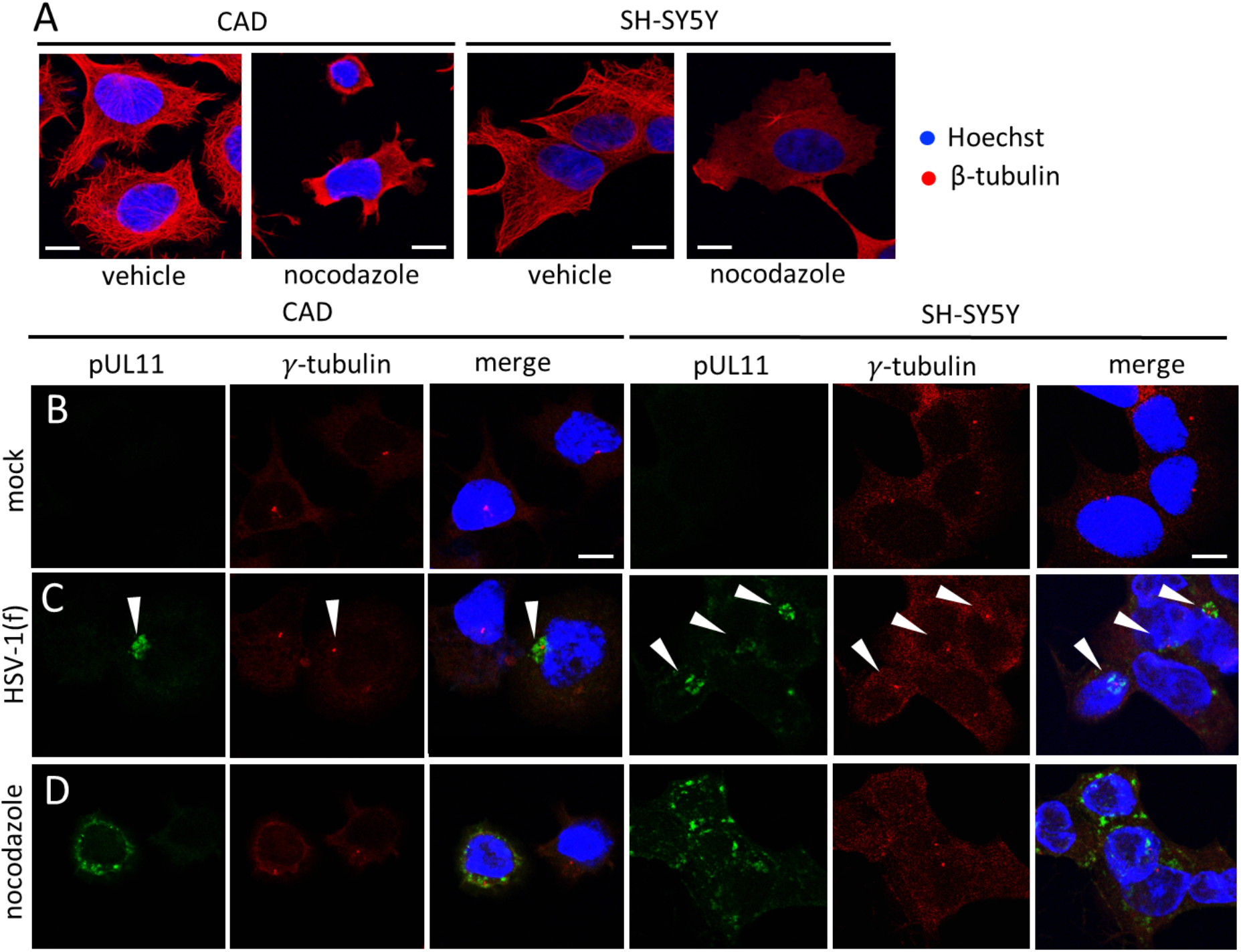
The formation of HSV cVAC is microtubule-dependent. (A) CAD and SH-SY5Y cells were treated with either DMSO (upper panel) or 5μg/ml of nocodazole (lower panel) for 6 hs before fixation and stained for β-tubulin (red). Scale bars represent 10 μm. (B-D) CAD and SH-SY5Y cells were stained for pUL11 (green) and γ-tubulin (red). (B) Mock-infected cells were treated with DMSO. (C) Cells were infected with WT HSV-1(F) and treated with DMSO as described in Figure 8. White arrowheads indicate HSV cVACs. (D) Cells were infected with WT HSV-1(F) and treated with nocodazole as described in Figure 8.

**Figure 8.**
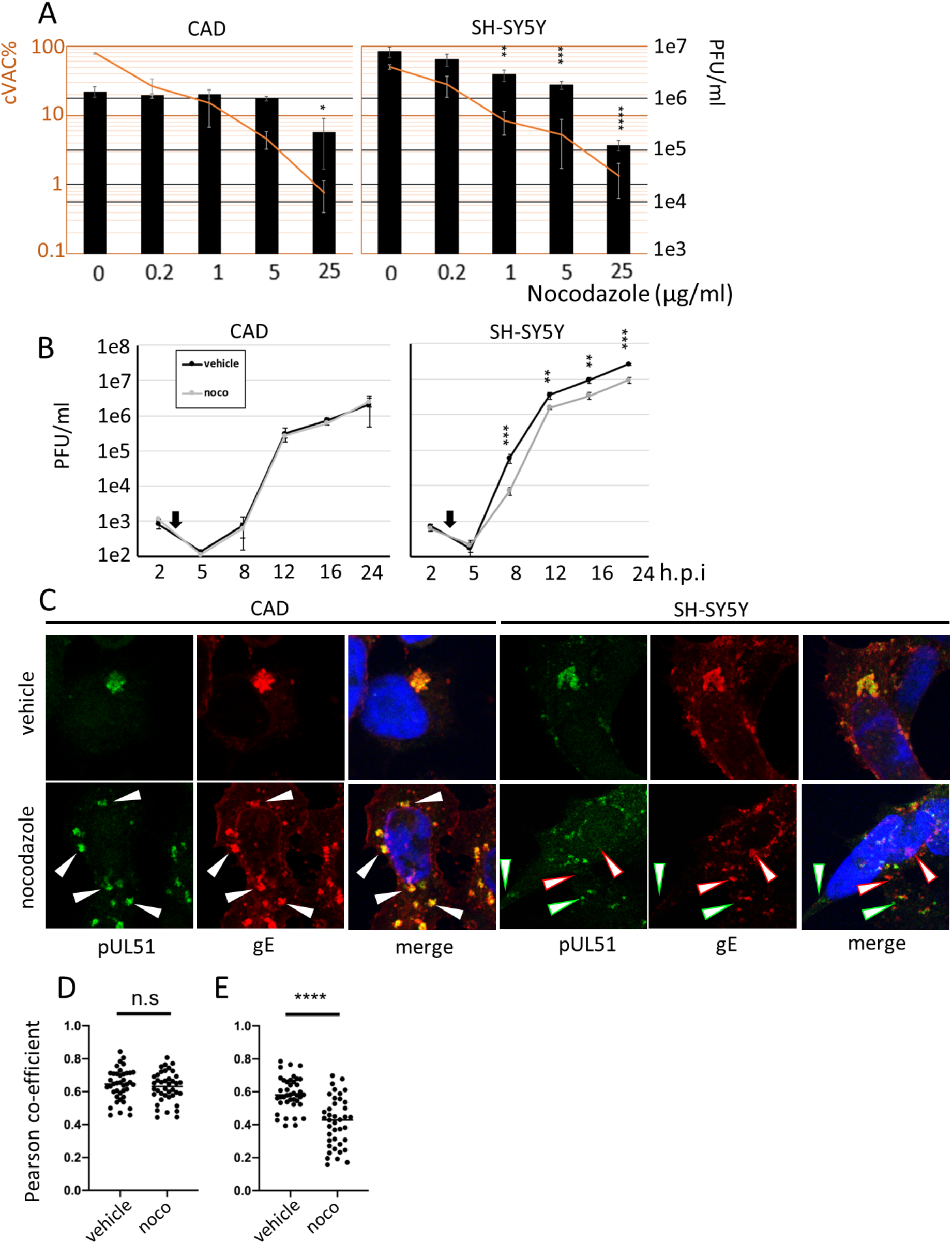
Nocodazole has cell type-specific effects on viral production. (A) The percentage of infected cells that form the cVAC decreases in respond to increasing concentration of nocodazole treatment in infected cells, with the viral production also decreases proportionally in infected SH-SY5Y cells. Orange lines present percentage of infected cells that form a unitary cVAC. Each cVAC is defined by pUL11 concentration around the γ-tubulin-defined centrosome. Infected cells were fixed at 12 h.p.i. Error bars represent standard deviation of three independent experiments. Black columns indicate viral production determined by single step growth (SSG) at 16 h.p.i as descried in method and material. (B) Growth curve of WT HSV-1(F) on DMSO (black line) or nocodazole (gray line) treated cells. Cells were infected as in (A). The infection medium was replaced with medium containing either DMSO or nocodazole at 3 h.p.i. (black arrows). (C) Cells were treated, infected, and fixed the same as in (A), except they were stained for pUL51 (green) and gE (red). For nocodazole treated CAD cells, white arrowheads indicate periphery puncta that are positive for both pUL51 and gE. For nocodazole treated SH-SY5Y cells, green arrowheads indicate pUL51-only puncta and red arrowheads indicate gE-only puncta. (D-E) Quantification of pUL51-gE co-localization in cells in (C). Each dot represents the Pearson co-efficient of pUL51 and gE in a random infected cell. For all statistics, n.s.: P>0.05; *: P<0.05; **: P<0.01; ***: P<0.005; ****: P<0.001.

### Nocodazole causes cell type-specific growth defect

Since nocodazole profoundly disrupts HSV cVAC formation, its net consequence, namely viral production, was investigated by single-step growth (SSG) assays. As shown in Figure 8A, viral progeny production was reduced in SH-SY5Y cells in a dose-dependent manner that proportionally correlates with the reduction in cVAC formation, but the same effect was not observed in CAD cells. This trend remained unchanged as late as 24 h.p.i (Figure 8B).These results suggest that nocodazole treatment is unable to change certain aspects of the HSV assembly compartment in CAD cells even if this compartment is no longer concentrated at one single region in the cell. Thus, we hypothesized that in nocodazole treated CAD cells, HSV assembly compartment is still able to form as individual periphery sites, similar to it in epithelial cells. To test this hypothesis, nocodazole-treated HSV-infected CAD and SH-SY5Y cells were stained for viral protein pUL51 and gE. As expected, pUL51 and gE co-localization remained unchanged in nocodazole treated CAD cells but show significant reduction in nocodazole treated SH-SY5Y cells (Figure 8, C and D).

### Microtubule-associated motors also play a role in cVAC maintenance

To further assess the contribution of microtubule components in HSV cVAC formation, we selectively disrupted the function of dynein, a microtubule-associated motor, with a dynein motor ATPase catalytic site inhibitor, ciliobrevin D (Firestone et al., 2012). Uninfected CAD and SH-SY5Y cells were treated with ciliobrevin D and stained for Rab11 as an indicator of dynein motor function. With 200μM of ciliobrevin D, which was the highest concentration tested, dispersal of Rab11 puncta was only observed in CAD cells (Figure 9A). To test whether ciliobrevin D disrupts cVAC formation, CAD cells were synchronously infected with HSV-(F) and treated with ciliobrevin D at 3 h.p.i before staining for pUL11 and γ-tubulin. As expected, HSV cVAC can be disrupted in a dose-dependent manner (Figure 9, B and C), supporting the hypothesis that the HSV cVAC maintenance requires microtubule associated components.

**Figure 9.**
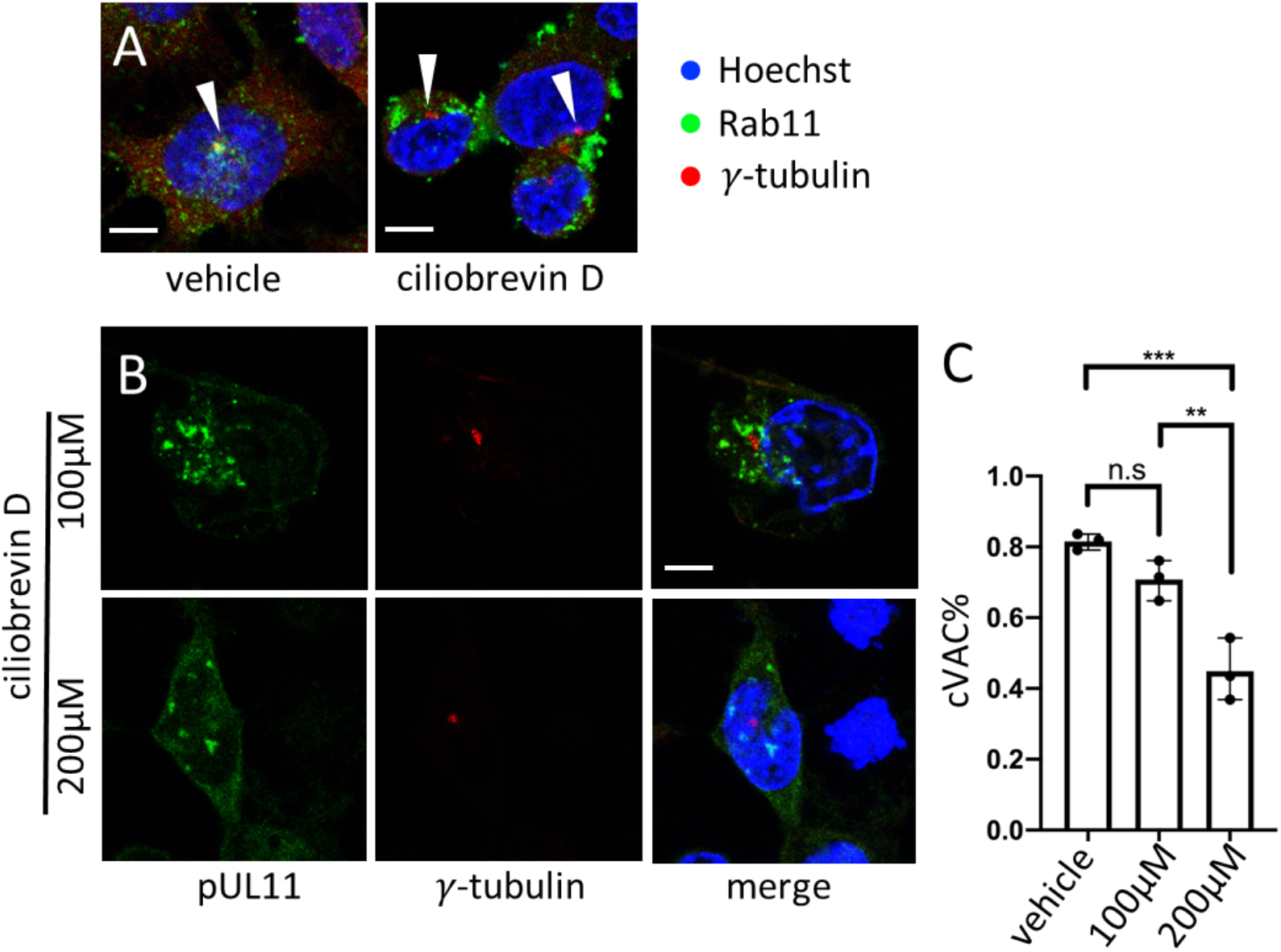
Dynein motors might also contribute to the maintenance of HSV cVAC. (A) CAD cells were treated with DMSO or 200 μM ciliobrevin D for 6 hs before fixation and staining for Rab11 (green) and γ-tubulin (red). White arrowheads indicate centrosomes. Scale bars represent 10 μm. (B) CAD cells were infected and treated with DMSO or ciliobrevin D following the same protocol as in Figure 8. Cells were stained for pUL11 (green) and γ-tubulin (red). (C) Percentage of cells that form a unitary cVAC were counted as described in Figure 10. Error bars represent standard deviations of three independent experiments (n.s.: P>0.05; *: P<0.05; **: P<0.01; ***: P<0.005).

### HSV cVAC formation is not dependent on actin filaments

To rule out the possibility that the cVAC disruption in nocodazole or ciliobrevin D treatment is due to general collapse of the cytoskeleton, infected CAD and SH-SY5Y cells were then treated with cytochalasin B, an actin filament polymerization inhibitor (MacLean-Fletcher and Pollard, 1980). When treated with cytochalasin B, uninfected or infected cells round up similarly to nocodazole treated cells, however, HSV cVAC still forms at the same efficiency in infected cells (Figure S2, A-C), suggesting HSV cVAC formation is not actin dependent.

### pUL51 plays important role in the maintenance of HSV cVAC morphology

A well-established literature has described various viral proteins in HCMV that are essential for proper cVAC morphology and viral production, many of which have homologs with similar functions in HSV. HCMV kinase pUL97 and its HSV homolog pUL13 have important functions in viral nuclear egress (Kato et al., 2006, Cano-Monreal et al., 2009, Marschall et al., 2005, Milbradt et al., 2009, Sharma et al., 2015, DeRussy et al., 2016) and have been implied to play role in viral cytoplasmic assembly (Prichard et al., 2005, Goldberg et al., 2011, Eaton et al., 2014, Gershburg et al., 2015). It was shown that the loss of pUL97 in HCMV results in abnormal cVAC formation (Prichard et al., 2005), however, it was unclear whether it was due to the defect in nuclear egress or the defect in cytoplasmic assembly. To test whether the presence of capsid in the cytoplasm is required for HSV cVAC formation, CAD and SH-SY5Y cells were infected with an HSV virus lacking the expression of major capsid protein VP5. As shown in Figure 10D, the HSV cVAC still formed, suggesting viral capsid is not required. Since the more profound role of pUL97 in HCMV and pUL13 in HSV is nuclear lamin network disruption (Kato et al., 2006, Sharma et al., 2015), it is also possible that the viral-induced nuclear membrane alteration is required for the formation of HSV cVAC. To test this hypothesis, CAD and SH-SY5Y cells were infected with an HSV virus lacking the expression of pUL34, a subunit of HSV nuclear egress complex (NEC) (Reynolds et al., 2001). Since pUL34 is a protein that is essential for lamin A/C phosphorylation (Bjerke and Roller, 2006), emerin phosphorylation (Leach et al., 2007), and nuclear shape alteration (Vu et al., 2016, Bjerke and Roller, 2006), a UL34-null virus should have severe defects in most major pathways of viral-induced nuclear membrane change. As shown in Figure 10E, loss of pUL34 did not disrupt the formation of HSV cVAC, suggesting lamin A/C and emerin phosphorylation are not essential for HSV cVAC formation.

HCMV tegument protein pUL71 was shown to be important for the cVAC formation and viral assembly (Schauflinger et al., 2011, Meissner et al., 2012, Womack and Shenk, 2010). Also, the N-terminus of pUL71 forms a tyrosine-based retrieval motif (Yxxφ), which upon mutation results in the accumulation of pUL71 on the cytoplasmic membrane, in addition to viral assembly defect (Dietz et al., 2018). The HSV tegument protein pUL51, a homolog of HCMV pUL71, has a conserved N-terminal Yxxφ motif at residue 19-22 (Roller et al., 2014). Based on these, we hypothesize that the conserved retrieval motif in pUL51 is also essential for the maintenance of HSV cVAC. To test this hypothesis, CAD and SH-SY5Y cells were infected with a mutant HSV of which tyrosine reside in the hypothetical Yxxφ motif was replaced with an alanine (pUL51_Y19A_). As shown in Figure 10F, cVAC no longer formed in the HSV mutant infected cells. In addition, pUL51 no longer efficiently co-localized with glycoprotein gE (Figure 10, G and H), suggesting that not only the concentrated HSV cVAC, but also periphery assembly sites were disrupted by the pUL51_Y19A_ mutation. Unsurprisingly, the pUL51_Y19A_ mutation also causes viral growth defect in both CAD and SH-SY5Y cells (Figure 10, I and J), which is consistent with our hypotheses that (i) viral assembly sites formation is important for efficient viral production, and (ii) pUL51 plays essential role in HSV cVAC formation.

**Figure 10.**
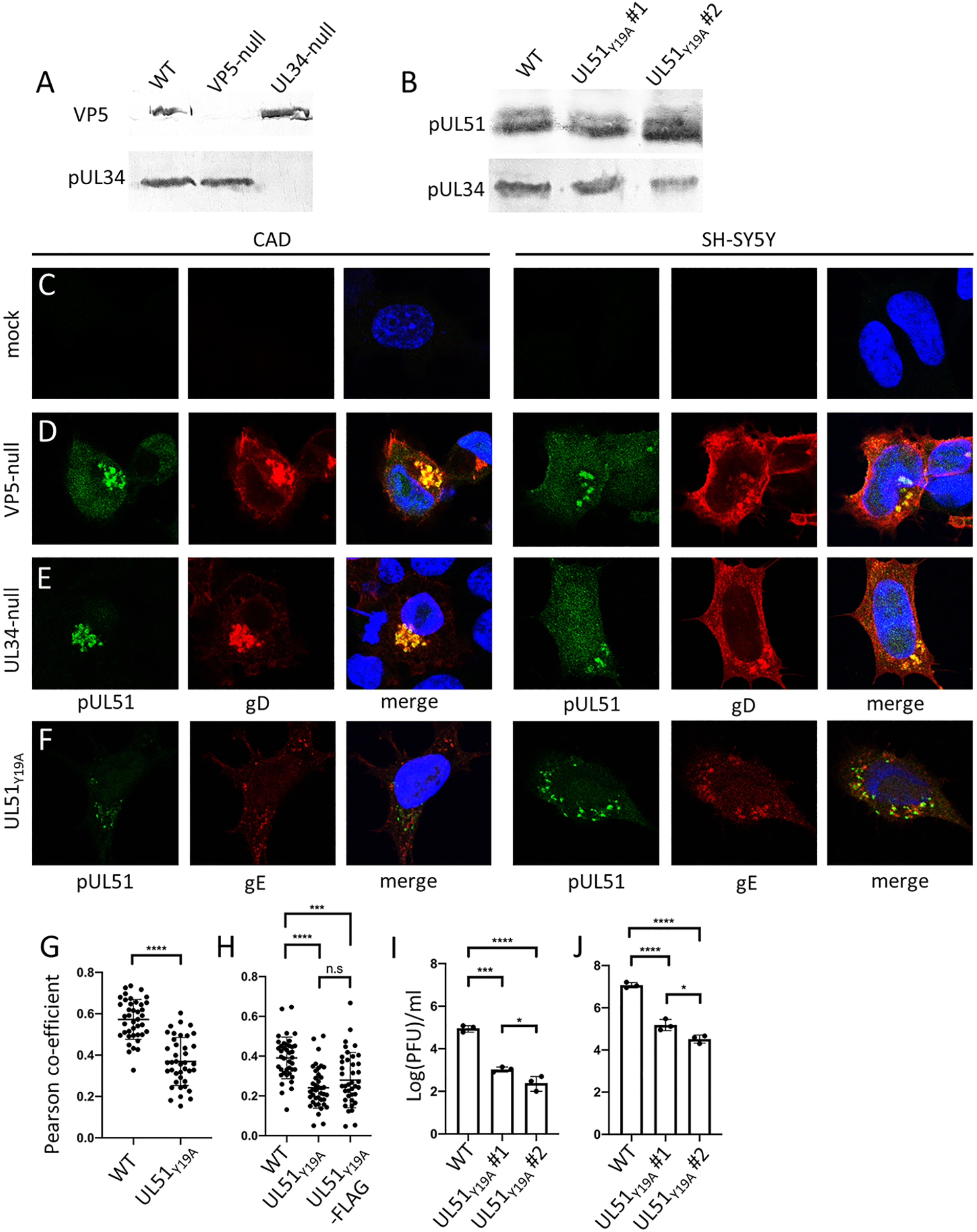
Viral structure proteins that are important in the maintenance of HSV cVAC. (A-B) Western blot of HSV BAC-derived viruses-infected cells. Lysate of infected SH-SY5Y cells (m.o.i.=5) was harvested 16 h.p.i. (C-F) CAD and SH-SY5Y cells were infected with HSV-1 mutant viruses (m.o.i=5) and stained for indicated viral proteins at 12 h.p.i. (G-H) Quantification of pUL51-gE colocalization in WT or mutant HSV-1 BAC viruses-infected CAD (G) or SH-SY5Y (H) cells by Pearson co-efficient. (I-J) SSG of indicated HSV-1 BAC-derived viruses on CAD (I) or SH-SY5Y (J) cells. For all statistics, n.s.: P>0.05; *: P<0.05; **: P<0.01; ***: P<0.005.

## Discussion

We have described an HSV infection-induced structure that concentrates at least six viral structural proteins specifically in neuronal cells. While we have only stained for only six viral structural proteins, they represent core components of at least two viral membrane protein complexes, including the fusion complex that is composed of gD, gB and gH/L (Rodger et al., 2001) and a cell-cell spread specific complex gE-gI (Johnson and Feenstra, 1987). All of these proteins have been reported to form stable complexes with other HSV structural proteins, and so it seems highly likely that other interacting viral factors including gM (Crump et al., 2004), gK/pUL20 (Lau and Crump, 2015), pUL7 (Roller and Feters, 2015), VP16 (Johnson et al., 1984), and VP22 (Farnsworth et al., 2007) also concentrate at the same structure. This is consistent with the hypothesis that the observed structure is a cVAC at which all of the components necessary for cytoplasmic envelopment are organized. Furthermore, almost all capsid envelopment events were observed concentrating to one region located to one side of the nucleus in CAD cells by TEM imaging (Figure 3). These capsids likely correspond to the capsid concentration in UL35-RFP virus infected cells (Figure 3), where is also where all viral proteins concentrate. Similar concentration of viral proteins and envelopment events have also been observed in other viruses that form cytoplasmic assembly centers, such as HCMV and reovirus (Tenorio et al., 2019).

While the viral structural proteins that we have tested all concentrate in the neuronal cVAC, none are exclusively localized to this structure, and they vary considerably in the proportion of the protein found there. pUL11 and pUL51 for example are highly concentrated in the cVAC (Figure 1). In contrast, gD, while concentrated in the cVAC is easily detectable on the cell surface and on membranes throughout the cytoplasm (Figure 1). It is unsurprising that all of the structural proteins have some presence outside of the cVAC, since they must be present in fully assembled egressing virions and as biosynthetic intermediates such as integral membrane proteins in the ER. Furthermore, several of the virion envelope glycoproteins contain plasma membrane retrieval motifs in their cytoplasmic tails and are thought to be retrieved from the plasma membrane and arrive at the site of cytoplasmic envelopment through endosomal trafficking (Alconada et al., 1999, Beitia Ortiz de Zarate et al., 2007, Crump et al., 2004). Interestingly, gD, which is the most widely distributed of the proteins we tested, does not have a retrieval motif (Crump et al., 2004) suggesting that possession of such a motif might be related to the mechanism of trafficking to the cVAC.

It is possible that concentrations of viral structural proteins could also reflect formation of aggresomes or autophagosomes, and be unrelated to virus assembly events. If so, concentration of viral proteins would be accompanied by recruitment of markers of autophagosomes or lysosomes. However, the autophagy marker LC3 does not concentrate to the cVAC (Figure 4), suggesting these membranes are not canonical autophagosomes. Also, the lysosomal marker LAMP1 does not co-localize with viral protein puncta (Figure 4), suggesting the cVAC is not a degradation compartment that involves lysosome. Combining these two observations with the stability of the cVAC throughout the infection (Figure 1F), we conclude that the cVAC is not a canonical degradation compartment. We did observe partial and transient co-localization of LAMP1, with virus structural proteins early in infection in some cells. This might reflect an early host antiviral response that inhibits virus production, or it may represent a virus-induced alteration in lysosome organization to promote replication.

The formation of a unique cVAC has not previously been reported in HSV infection, but appears to be a constant feature of infection of neuronal cells, as we have observed the cVAC structure in at least two cancerous neuronal cell lines, one of murine origin and one of human origin, and in primary mouse neurons. While it would be of great interest to observe whether a cVAC structure forms in human trigeminal neurons during a natural oral infection as well, such clinical samples would be rare and it is not known how large a subset of these neurons undergo lytic infection (Held and Derfuss, 2011).

Since a single HSV cVAC does not form in infected epithelial cells, and since, within the mixed culture of primary neural cells, only neurons, but not astrocytes, allow the formation a cVAC (Figure 2), it is likely that formation of a single cVAC is a specific viral adaptation for infection of neurons. Also, despite that HSV forms cVAC in both CAD and SH-SY5Y cells, there exists considerable variations in their morphology (Figure 1), which suggests the importance of host factors that may vary between human and mouse species or between different neuronal subsets. One factor that might have contributed to the difference between HSV assembly compartment in neurons and epithelial cells is cell polarity. In epithelial cells, cell polarity follows the apical-basolateral axis, while the cell polarity in neurons follows the axon (Bartolini and Gundersen, 2006, Bonifacino, 2014). Since HSV capsids and vesicles that are positive for viral structural proteins move along the microtubule network (Kratchmarov et al., 2012), it is possible that these viral components, despite being transported in the same manner along microtubules, are trafficked to distinct compartments, such as basolateral side in epithelial cells and centralized compartment at cell body in neurons, due to specific cell polarity. Similarly, another possible source of HSV assembly compartment difference is the availability of host microtubule components in neurons. This includes as least three non-mutually exclusive aspects: (i) the lack of factors for non-centrosomal microtubule nucleation site in neurons. It was described that Golgi membrane can anchor microtubule minus end and serve as a MTOC in several types of somatic cells (Sanchez and Feldman, 2017), and indeed, in HCMV-infected cells, microtubules preferentially polymerize from Golgi-derived membranes instead of the centrosome (Procter et al., 2018). Since there has been no evidence hitherto suggesting that such alternative MTOC sites can occur efficiently in mammalian neurons (Sanchez and Feldman, 2017), it is possible that the deficiency or absence of alternative microtubule nucleation factor necessitates the centrosome as the viral assembly site in neurons, hence the unitary cVAC. (ii) Different expression level or distribution of certain dynein components or cargo adaptors. Higher eukaryotes encode several isoforms of dynein intermediate and light chains, which are important for cargo specificity (Pfister et al., 2006, Dixit et al., 2008). Also, is has been described that the same trafficking motif of the same protein might utilize different adaptor protein subunits in neurons and epithelial cells (Bonifacino, 2014, Farias et al., 2012). It is possible that viral tegument and glycoproteins can only utilize a certain subset of these dynein motors or cargo adaptors, which in neurons only provides trafficking towards the centrosome while in epithelial cells provides diverse trafficking destinations. (iii) The centrosome in neuronal cells is resistance to viral manipulation. It has been observed that in HSV-infected epithelial cell, centrosome disappears in many cells (Pasdeloup et al., 2013), while in most infected neuronal cells, the centrosome appears to be intact (Figure 3, 7, 9, S2). It is possible that dispersed assembly compartments of HSV in epithelial cells is caused by the loss of centrosome, while in neurons, the stability of the centrosome allows it to out-compete alternative sites and to serve as the major MTOC, which naturally concentrates all viral structural proteins.

### The organization of host membranes

The identity of the membrane at which HSV cytoplasmic envelopment occurs in epithelial cells is currently controversial, with evidence supporting both trans Golgi and endosomal membranes, but not ER (Lv et al., 2019). The arrangement of host Golgi, ER, and some endosomal membranes in the HSV neuronal cVAC is broadly consistent with previous results derived from epithelial cell studies. ER membranes are not included in the neuronal cVAC (Figure S1). In contrast, cis- and trans-Golgi-derived membranes concentrate to the cVAC but do not completely co-localize with viral structural proteins (Figure 5). This is similar to the previously described localization pattern of Golgi-derived membranes in HSV-infected epithelial cells, in which viral capsid and tegument proteins localize adjacent to, and sometimes overlapping with Golgi marker staining (Sugimoto et al., 2008, Hollinshead et al., 2012). Hollinshead et al., have shown that some HSV in epithelial cells undergoes envelopment at membranes that are derived from endosomes (Hollinshead et al., 2012). Consistent with this idea, we observed concentration of Rab11 endosomes in the cVAC with extensive co-localization between Rab11 and virus structural proteins (Figure 6). Hollinshead and colleagues also showed that Rab5 and Rab11 are important in HSV assembly in HeLa cells (Hollinshead et al., 2012). Rab11 is a recycling endosome marker that has been proposed to play role in the export of HSV cytoplasmic virions (Spearman, 2018, Bucci et al., 2014). While concentration of Rab11 at the neuronal cVAC is consistent with a role in virion export, the extensive co-localization between Rab 11 and the viral structural proteins raises the possibility that Rab11 endosomes may be an important site of envelopment.

Rab5 is an early endosome marker that, in neurons, retrieves vesicles from both cytoplasmic membrane of the cell body and synaptic membranes from the distal end of axon (Spearman, 2018, Bucci et al., 2014). We have observed that Rab5 does not concentrate in the HSV neuronal cVAC, and instead is localized in peripheral puncta, where it co-localizes with some of the cytoplasmic gE (Figure 6C). It is possible that these puncta represent retrieval vesicles that contain viral glycoproteins being trafficked to the cVAC.

### Species specific requirement of microtubule network

The HSV cVAC in cancerous neuronal cells forms around the MTOC defined by γ-tubulin staining (Figure 3, 7), suggesting that it is formed and maintained by retrograde transport of membranes and proteins toward the MTOC by microtubule-associated motors. Consistent with this hypothesis, both nocodazole, a microtubule polymerization inhibitor (De Brabander et al., 1976), and ciliobrevin D, a dynein motor ATPase inhibitor (Firestone et al., 2012), are able to disrupt cVAC morphology in CAD cells (Figure 6–9). However, the requirement for an intact microtubule network and, therefore, for a normally formed cVAC for viral assembly in neuronal cell lines is cell-type dependent. Nocodazole treatment affects virus production in a dose-dependent manner in SH-SY5Y, but not in CAD cells. This suggests that in CAD cells, despite the dispersal of unitary cVAC around the MTOC, viral proteins are still able to associate with each other to form dispersed, peripheral assembly sites, and indeed we still observed co-localization of the viral structural proteins (Figure 8C). In contrast, in SH-SY5Y cells, tegument proteins and glycoproteins no longer co-localize with each other efficiently when microtubule network is disrupted (Figure 8C). The mechanistic basis for this discrepancy is unclear, but possibilities include: (i) different sensitivities of the microtubule network to nocodazole. It has been described that herpesviruses can stabilize microtubules via acetylation or de-tyrosination (Wloga et al., 2017, Procter et al., 2018, Naghavi et al., 2013). Thus, a subset of microtubules in CAD cells might be resistant to nocodazole during HSV infection and still promote organization of peripheral assembly sites. (ii) Biosynthetic and retrieval pathways for membrane glycoproteins might differ between these cell types such that localization of assembly proteins to the same membrane compartment might occur without a need for microtubule-dependent trafficking in CAD cells. It is interesting that disruption of the cVAC by nocodazole in CAD cells creates a dispersed assembly apparatus that is morphologically similar to that observed in epithelial cells, and that virus assembly in epithelial cells is not sensitive to nocodazole treatment. Whether the similarities between the assembly compartments in nocodazole-treated CAD cells and untreated epithelial cells are merely superficial or reflective of similar mechanisms of assembly compartment formation is not clear.

### Similarities and discrepancies between HCMV cVAC and HSV cVAC

It is well-documented that HCMV forms a unitary cVAC in infected cultured fibroblasts (Alwine, 2012) and forms a dense cytoplasmic inclusion in cells from clinical samples (Gilloteaux and Nassiri, 2000). There are many similarities between the cVAC formed by two herpesviruses regarding host membrane arrangement, host cytoskeleton involvement, and viral factor requirement. First, we have observed that Golgi-derived membranes concentrate to the HSV cVAC, similar to the Golgi marker arrangement in HCMV-infected cells (Das et al., 2007, Das and Pellett, 2011), suggesting herpesvirus conserved requirement for Golgi function. On the other hand, the localization pattern of ER membranes in HSV-infected neuronal cells (Figure S1) is similar to the common laboratory strains of HCMV-infected fibroblasts, in which ER membranes mostly retain their natural structure (Zhang et al., 2019). This is different from the ER morphology of wild-type HCMV strain TB40/E-infected cells, in which ER membranes are deformed by dilation and ruffling (Zhang et al., 2019). This strain-specific difference is due to the quick loss of a single gene UL148 (which encodes an ER-modification protein in HCMV strain TB40/E) during the passage of the virus in cell cultures (Zhang et al., 2019). UL148 has no homolog in HSV, thus the resemblance between ER patterns in HSV-infected cells and HCMV-infected cells is not surprising.

Other host membrane markers, notably Rab family GTPases, localize to the cVAC of HSV and HCMV in patterns that are consistent with their roles in the vesicle trafficking and in viral assembly. For example, Rab11 participates in the viral assembly of both HSV and HCMV (Hook et al., 2014, Krzyzaniak et al., 2009), and indeed, Rab11 concentrates to the cVAC of both viruses (Das and Pellett, 2011, Das et al., 2007)(Figure 6, I and M). On the other hand, Rab5 is involved in HSV assembly, thus we observed its co-localization with periphery gE puncta (Figure 6C); while Rab5 is unlikely to support HCMV assembly as it is not present in the virion (Spearman, 2018), thus no particular co-localization between HCMV viral proteins and Rab5 was observed in infected cells (Das et al., 2007).

The requirement for cytoskeleton components is also similar between HCMV and HSV. In cells infected with either virus, nocodazole is able to disrupt cVAC morphology (Indran et al., 2010)(Figure 7, 8), suggesting that utilization of the microtubule network is a conserved mechanism for herpesvirus assembly. Also, sensitivity of the HSV cVAC to ciliobrevin D suggests that dynein motors are important for cVAC maintenance (Figure 9), as discussed above, however the potential off-target effect of the dynein ATPase inhibitor cannot be excluded. Inhibition of cVAC formation or maintenance by expression of the dynactin subunit p150Glued CC1 domain, which is a dominant negative inhibitor of dynein motor function, has been demonstrated in HCMV-infected cells (Buchkovich et al., 2010). However, the much shorter lifecycle, and the dynein dependence of early events in infection, make this approach more difficult for HSV. Conversely, small molecule inhibitors like nocodazole and ciliobrevin D also have limitations in assessing the consequences of microtubule disruption in HCMV-infected cells, as cell viability dwindles quickly after treatment.

We hypothesize that homologous viral proteins might also function similarly in cells infected with HCMV and HSV. pUL71 of HCMV, which is a homolog of HSV pUL51, was shown to be important for viral cytoplasmic envelopment and proper cVAC morphology (Schauflinger et al., 2011). Reciprocally, in this study, we have shown that pUL51 in HSV is also important for cVAC maintenance, as mutations in HSV pUL51 disrupts cVAC formation (Figure 10F). Interestingly, although the hypothetical tyrosine-based trafficking motif in the N-terminus of pUL51 is conserved in several herpesviruses and has been shown to be a cytoplasmic retrieval motif in HCMV homolog pUL71(Dietz et al., 2018), it is as yet unclear whether the same holds true for HSV. It is possible that the tyrosine residue at position 19 in pUL51 is indeed a trafficking motif, which upon mutation prevents pUL51 from concentrating to the HSV cVAC and, in turns, causes failure in cVAC formation.

### cVAC as a model for HSV assembly study

The unitary HSV cVAC in neuronal cells provides great advantages in the study of HSV cytoplasmic assembly, as it allows accessing of phenotypes of mutant viruses that show no obvious morphology change under microscope in epithelial cells, which in turn can be used to answer difficult yet fundamental questions about HSV biology. One of the outstanding questions in HSV assembly is that whether tegument and glycoproteins are pre-deposited onto a membranous structure that serves as the budding site, or it is the capsid itself recruits vesicles that contain various viral structural proteins. In this study, we have shown that the formation of the viral assembly compartment is independent of capsid and capsid nuclear egress (Figure 10D, E), confirming a tegument- and glycoprotein-centric model for organization of HSV assembly centers. Further supporting this model, a mutation in tegument protein pUL51 was able to not only disperse the centralized cVAC, but also disrupt co-localization between viral components (Figure 10F), suggesting tegument proteins might be essential for the maintenance of the HSV assembly compartment. All in all, we believe that the cultured neuronal cells will be powerful tools to study HSV assembly.

## Materials and Methods

### Cell cultures

CAD cells (gift from David Johnson at Oregon Health & Science University) were maintained in Dulbecco modified Eagle medium (DMEM) F-12 (Thermo Fisher Scientific) supplemented with 10% fetal bovine serum (FBS) and penicillin-streptomycin (P/S). SH-SY5Y cells (gift from Stevens Lewis at the University of Iowa) were maintained in Opti-MEM supplemented with sodium pyruvate, non-essential amino acid, P/S, and 10% FBS. Primary mouse cerebral cortex neurons and mouse glial feeder cells in primary culture were prepared by a method described previously (Mitchell et al., 2018) with slight modifications. In brief, the cerebral cortex was dissected from wild-type mice at postnatal days 0-1. Tissues of cerebral cortex were taken from the region immediately dorsal to the striatum. The dissected tissues were trypsinized and mechanically dissociated. The cells were plated on 12-mm coverslips (thickness No. 0, Carolina Biological Supply, Burlington, NC) in 24-well plates, at the density of 10,000 cells per coverslip. The coverslips had previously been seeded with an astrocytic feeder layer at the density of 20,000 cells per coverslip for 7-9 days, prepared from the CA3-CA1 region of the wild-type mouse hippocampus at postnatal days 0-1. The cultured neurons were infected with the virus after 3 days *in vitro* (DIV). All cells were cultured in a humidified incubator at 37°C, with 5% CO_2_.

Animal care and use procedures were approved by the University of Iowa’s Institutional Animal Care and Use Committee, and performed in accordance with the standards set by the PHS Policy and The Guide for the Care and Use of Laboratory Animals (NRC Publications) revised 2011. Every effort was made to minimize suffering of the animals.

### Plasmid and DNA transfection

pcDNA-GFP plasmid was purchased from Addgene (Plasmid #13031). CAD or SH-SY5Y cells were seeded into 24-well tissue culture plates containing glass coverslips at 1×10^5^ cells per well 17-24 hs before transfection. Cells were then transfected with X-tremeGENE™ HP DNA Transfection Reagent (Millipore-Sigma) or Lipofectamine^®^ Reagent (Invitrogen) following product protocols.

### Construction and production of recombinant viruses

The properties of HSV-1(F) virus have been described before (Ejercito et al., 1968). The HSV-1(F) BAC virus was a gift from Yasushi Kawaguchi at the University of Tokyo (Tanaka et al., 2003). The recombinant virus with a point mutation at residue 19 in UL51 was constructed using this HSV-1(F) BAC as previously described (Roller et al., 2010). The mutagenesis insertion DNA fragment was first amplified by PCR as two fragments: fragment one (fwd primer: 5’-GCCCCGAGGAACAAGCTGAGATGATCCGCGCGGCCGGCTCGGATCCTAGGGATAACAGGG-3’; rev primer 5’-CACTACGCGGCTGCTCAAACC-3’) and fragment two (fwd primer: 5’-GCGGTTGTTGGCGCTCTCG-3’; rev primer: 5’-GGCCGCGCGGATCATCTCAGCTTGTTCCTCGGGGCGCGCTCTCTAGAGGCCGCGGCGTTG-3’). These two fragments were assembled together by PCR with a pair of outer primers: fwd primer: 5’-CGGGGCTATATGTGGCTGGGGAGCGCGCCCCGAGGAACAAGCTGAGATG-3’; rev primer: 5’-CTCCGCCTCCGAGGGCGGAACGGCCGCGCGGATCATCTC-3’. Recombinant viruses were rescued by transfecting BAC DNA into Vero or Vero-derived complementing cells. Virus containing mutations in the UL51 gene sequence were amplified on UL51-complementing cells (Roller et al., 2014). HSV-1(F) BAC-derived virus containing deletions in the UL34 gene sequence have been described before (Roller et al., 2010) and were amplified on UL34-complementing cells (Roller et al., 2000). The HSV-1(F) VP5-null virus and VP5-complementing cells were kind gifts from Prashant Desai (Desai et al., 1993). HSV-1(F) UL35-RFP BAC virus was a gift from Cornel Fraefel at the University of Zürich (de Oliveira et al., 2008). HSV-2 strain R519-derived BAC virus (bHSV2-BAC38) was a gift from Kening Wang at National Institute of Allergy and Infectious Diseases (Wang et al., 2012).

### Viral infections

When infecting CAD and SH-SY5Y cells, HSV was diluted in the following infection medium: DMEM containing 1% heat-inactivated serum and P/S. In synchronized infection, cells were incubated in the infection medium containing HSV at 4°C for 1h before replacing with warm infection medium to allow viral entry. This medium change is designated as 0 h.p.i. For drug treatment, the infection medium was replaced with the same medium containing respective compound (or vehicle only). Stock solutions of nocodazole (AdipoGen), ciliobrevin D (EMD Millipore), and cytochalasin B (AdipoGen) were made in DMSO. For SSG and growth curve experiments, after 4°C incubation and medium replacement, monolayer of confluent cells were incubated at 37°C for 1 h to allow viral entry. Then the monolayer was incubated in a citrate buffer (50 mM sodium citrate, 4 mM KCl, adjusted to pH 3.0 with HCl) for 1 min to inactivate residual virus. The cell monolayer was then washed twice in PBS and incubated in infection medium before drug treatment at 3 h.p.i.

When infecting primary cortex neurons, HSV was diluted in the 1:1 mixture of plating medium and growth medium described above. Cultures were incubated in the inoculum for 1 h before replacing with the same media mixture.

Transduction of cells using the CellLight Golgi-GFP baculovirus vector (Thermo-Fisher) that expresses GFP-N-acetylgalactosaminyltransferase was performed according to the manufacturer’s instructions 24 hours prior to infection of cells with HSV-1.

### Indirect immunofluorescence and analysis

Transfected or infected cells on 13-mm-diameter coverslips in 24-well tissue culture plates were fixed with 3.7% formaldehyde for 20 minutes before washing with PBS two times. The immunofluorescence (IF) buffer used to dilute antibodies or antiserum is PBS that contains 1% triton X100, 0.5% sodium deoxycholate, 1% egg albumin, and 0.01% sodium azide. Fixed coverslips were first blocked with IF buffer containing 10% human serum before 1 to 2 hs-long primary antibody incubation (with 10% human serum) and 1 h secondary fluorescent antibody incubation. The primary antibodies reactive with viral proteins and cellular markers include 1:500 rabbit anti-UL51 antiserum (gift from Joel Baines), 1:500 rabbit anti-UL11 antiserum, 1:500 rabbit anti-UL16 antiserum, 1:500 rabbit anti-UL21 antiserum (all three provided by John Wills), 1:1,500 mouse [DL6] anti-gD monoclonal antibody, 1:500 mouse [L4] anti-gL monoclonal antibody (both gifts of Gary Cohen and Roselyn Eisenberg), 1:500 mouse monoclonal anti-gE antibody (Virusys), 1:500 mouse anti-MAP2 antibody (Sigma-Aldrich), 1:500 mouse anti-*γ*-tubulin antibody (Sigma-Aldrich), 1:500 rabbit anti-*γ*-tubulin antibody (Sigma-Aldrich), 1:500 mouse [H4A3] anti-LAMP1 antibody (DHSB), 1:250 rabbit anti-Rab5 antibody (Abcam), 1:250 rabbit anti-Rab11 antibody (Abcam), 1:500 mouse anti-GM130 antibody (BD Bioscience), 1:500 mouse polyclonal anti-Golgin97 antibody (Abcam), and 1:500 mouse anti-*γ*-tubulin antibody (Sigma-Aldrich). Actin filament was stained with 1:1,000 Alexa Flour™ 647 fluorescence conjugated phalloidin (Thermo Fisher Scientific). Nuclei were stained with 1:500 DNA dye Hoechst 33342 (Invitrogen). Stained coverslips were then mounted with ProlLong^™^ Diamond Antifade Mountant (Invitrogen) and allowed to cure in the dark at room temperature overnight. Confocal images were taken on a Leica DFC7000T confocal microscope equipped with Leica software. Signal patterns of interest were counted by eye. Co-localization efficiency of signals from different channels were measured with FIJI ImageJ software for each cell.

### Western blot

Cell lysates were separated on SDS-PAGE by size and transferred to nitrocellulose membranes before blocking in 5% nonfat milk plus 0.2% Tween 20 for at least 1 h. Then, membranes were probed with either 1:500 rabbit anti-UL51 antiserum, 1:500 mouse anti-VP5 antibody (Biodesign), or 1:250 chicken anti-UL34 antiserum (Reynolds et al., 2001) and visualized by alkaline phosphatase-conjugated secondary antibody.

### Transmission electron microscopy

Confluent monolayers of CAD cells on coverslips were infected with HSV-1(F) at a m.o.i of 5 for 12 h and then fixed by incubating in 2.5% glutaraldehyde in 0.1 M cacodylate buffer (pH 7.4) for at least 1 h. Cells were post-fixed in 1% osmium tetroxide, washed in cacodylate buffer, embedded in Spurr’s resin, and cut into 95-nm sections. Sections were mounted on grids, stained with uranyl acetate and lead citrate, and examined with a JEOL 1250 transmission electron microscope.

## Acknowledgments

The authors would like to thank the many investigators (listed individually in the methods section) for providing viruses, cell lines, and antibody reagents for these studies. We are also grateful to members of the Roller laboratory and to Keith Jarosinski and Wendy Maury for critical reading of the manuscript. This study was supported by NIH R21AI133155 and by the Carver College of Medicine, University of Iowa. H. K. and N. C. H. were supported by the University of Iowa.

---

**Figure S1.**
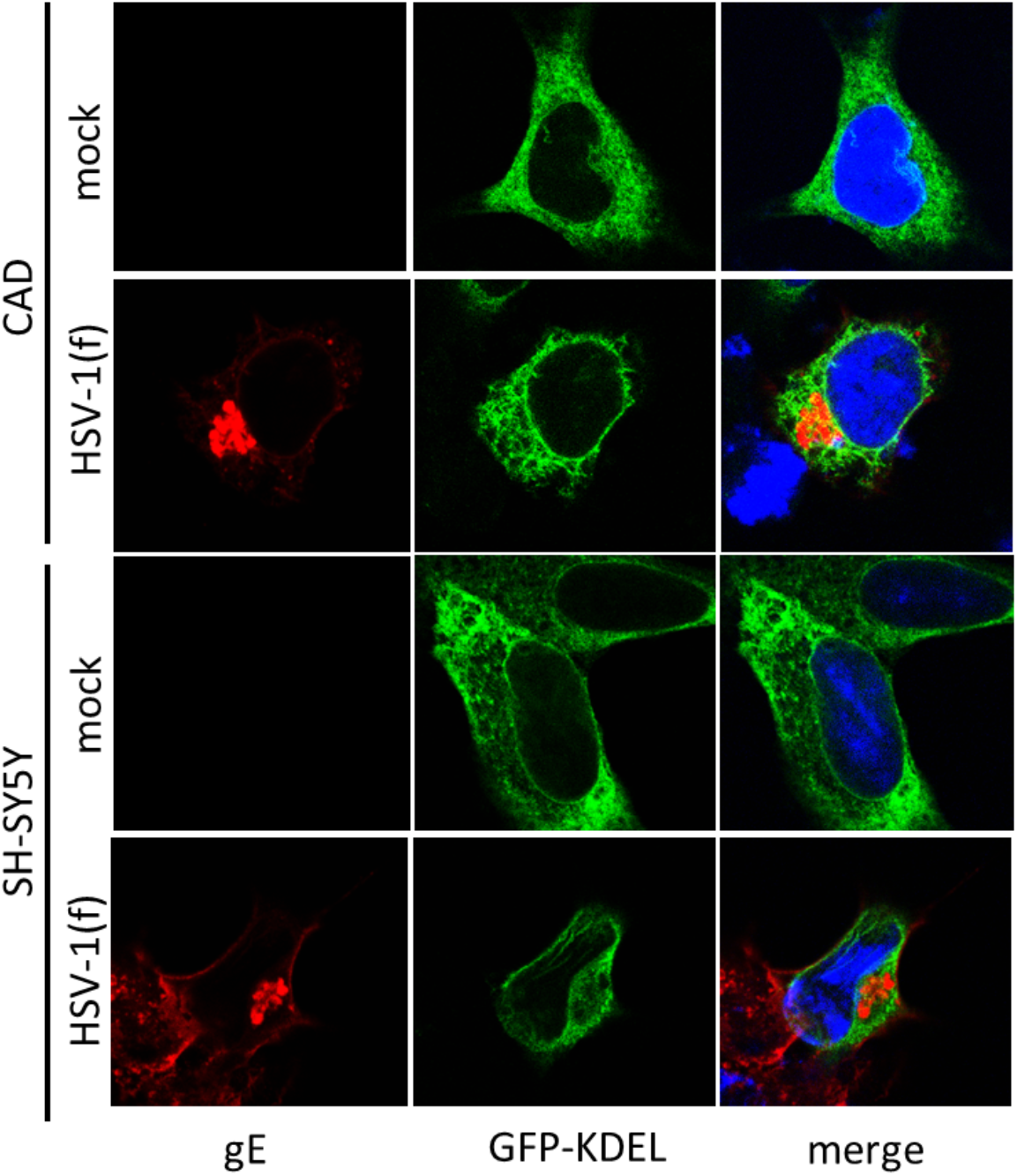
ER-derived membranes are not recruited to the HSV cVAC. CAD and SH-SY5Y cells were transfected with a GFP-KDEL expression plasmid at least 24 hs before mock or WT HSV-1(F) infection. Cells were infected as described in Figure 1 and stained for gE (red).

**Figure S2.**
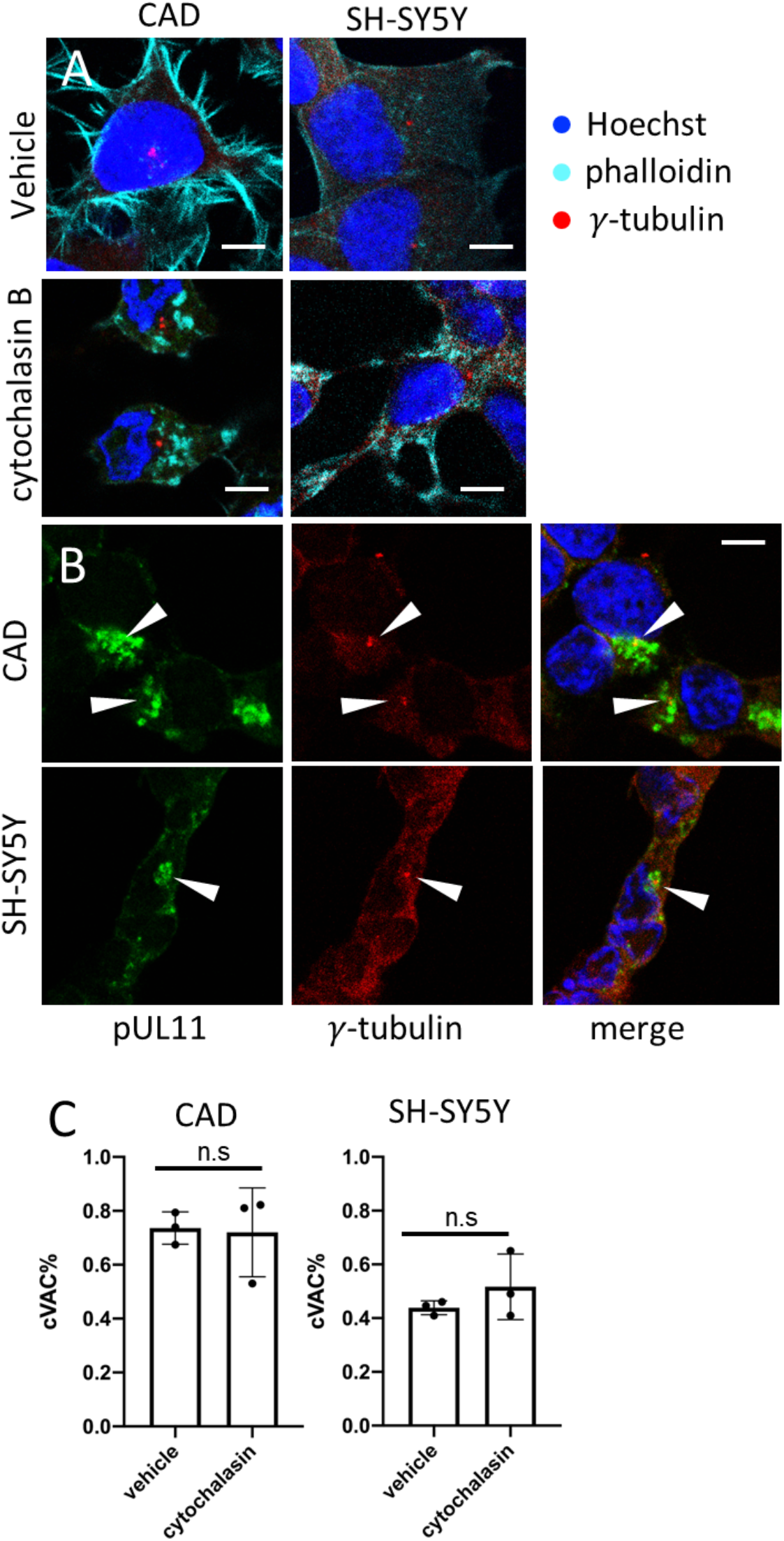
Actin filaments are not required to concentrate viral proteins to the HSV cVAC. (A) CAD and SH-SY5Y cells were treated with DMSO or cytochalasin B for 6 hs before fixation and staining for actin (cyan; stained with fluor-conjugated phalloidin) and γ-tubulin (red). Scale bars represent 10 μm. (B) Cells were infected and treated with DMSO or cytochalasin B following the same protocol as in Figure 8. Cells were stained for pUL11 (green) and γ-tubulin (red). White arrowheads indicate individual cVACs. (C) Percentage of cells that form a unitary cVAC were counted as described in Figure 10. Error bars represent standard deviations of three independent experiments (n.s.: P>0.05; *: P<0.05; **: P<0.01; ***: P<0.005).

